# Comparative proteomics of biofilm development in *Pseudoalteromonas tunicata* discovers a distinct family of Ca^2+^-dependent adhesins

**DOI:** 10.1101/2024.10.22.619756

**Authors:** Sura Ali, Alexander Stavropoulos, Benjamin Jenkins, Sadie Graves, Geoffrey Che, Jiujun Cheng, Huagang Tan, Xin Wei, Suhelen Egan, Josh D. Neufeld, Ulrich Eckhard, Trevor C. Charles, Andrew C. Doxey

## Abstract

The marine bacterium, *Pseudoalteromonas tunicata*, is a useful model for studying mechanisms of biofilm development due to its ability to colonize and form biofilms on a variety of marine and eukaryotic host-associated surfaces. However, the pathways responsible for *P. tunicata* biofilm formation are still incompletely understood, in part due to a lack of functional information for a large proportion of its proteome. Here, we used comparative shotgun proteomics to examine *P. tunicata* biofilm development throughout the planktonic phase to three stages of biofilm development at 24, 48, and 72 h. Proteomic analysis identified 232 proteins that were up-regulated during different stages of biofilm development, including many hypothetical proteins as well as proteins known to be important for *P. tunicata* biofilm development such as the autocidal enzyme AlpP, violacein proteins, S-layer protein SLR4, and various pili proteins. We further investigated the top identified biofilm-associated protein (Bap), a previously uncharacterized 1600 amino acid protein (EAR30327), which we designated as “BapP”. Based on AlphaFold modeling and genomic context analysis, we predicted BapP as a distinct Ca^2+^-dependent biofilm adhesin. Consistent with this prediction, a Δ*bapP* knockout mutant was defective in forming both pellicle and surface-associated biofilms, which was rescued by re-insertion of *bapP* into the genome. Similar to mechanisms of RTX adhesins, BapP-mediated biofilm formation was influenced by Ca^2+^ levels, and BapP is likely exported by a type 1 secretion system. Ultimately, our work not only provides a useful proteomic dataset for studying biofilm development in an ecologically relevant organism, but it also adds to our knowledge of bacterial adhesin diversity, emphasizing Bap-like proteins as widespread determinants of biofilm formation in bacteria.

## INTRODUCTION

Understanding the molecular processes that control biofilm development by environmental and host-associated microorganisms is a fundamental area of microbiology, with important industrial and ecological applications. Members of the *Pseudoalteromonas* genus (class *Gammaproteobacteria*) are commonly found in marine environments in association with biological surfaces and diverse eukaryotic hosts and play important roles in the ecology of marine ecosystems (1–3). One of the best studied species within this group is *P. tunicata,* a heterotrophic, Gram-negative bacterium first isolated from the tunicate, *Ciona intestinalis* (4). *P. tunicata* is also known to colonize algal host surfaces, sea-water biofilm communities, and likely other yet-to-be-identified living surfaces and host organisms in the marine environment (5, 6). Among *P. tunicata*’s characteristics is its ability to colonize and outcompete other species in natural biofilms (6, 7), which is in part due to its broad repertoire of antimicrobial capabilities (3–5, 8). Characterizing the molecular basis of *P. tunicata*’s biofilm formation, colonization, and antifouling activity is important not only in the context of understanding marine biofilm ecology (6, 8–10), but it could also reveal new biotechnological strategies for preventing harmful biofilm formation (e.g., in industrial settings or infections) (11, 12).

Several previous studies have investigated biofilm development in *P. tunicata* (6–8), which shares several general mechanisms with that of other biofilm-forming bacteria. The first step is the initial attachment/adhesion to surfaces (8). Typically, these initial interactions involve weak and reversible binding to a surface substrate, using adhesive structures such as flagella and pili (13). The genome of *P. tunicata* encodes a variety of flagellar genes including a proteolytically active variant of flagellin that might be involved in flagella-mediated interactions with surfaces or host cells (14, 15). In addition, the *P. tunicata* genome encodes a diversity of surface adhesion-related proteins capable of binding diverse substrates in the marine environment. These include type IV pili, curli, MSHA-like pili, capsular polysaccharide, chitin and cellulose-binding proteins, as well as specialized proteins for binding to extracellular matrix (ECM) components abundant in eukaryotic host surfaces (5, 16, 17). Among these is LipL32, which facilitates adhesion of *P. tunicata* to the ECM of its *Ciona intestinalis* host (18).

The transition of *P. tunicata* to a surface-associated lifestyle involves dynamic changes in its transcriptome and proteome (16). These changes include the production of anti-biofouling agents (e.g., antilarval, antibacterial, antialgal, and antifungal molecules) (3, 19), as well as the formation of the extracellular biofilm matrix that provides mechanical and protective scaffold for the biofilm. The ToxR-like transcriptional regulator WmpR is a key regulatory protein that controls stationary phase expression of antifouling inhibitors in *P. tunicata* (20). WmpR also controls the production of other bioactive compounds and pigments (20), and other proteins associated with adaptation to a biofilm lifestyle such as iron acquisition genes and type IV pili (8, 19). One of the key proteins produced by *P. tunicata* is the autocidal enzyme, AlpP, that causes cell lysis within biofilms (8). AlpP is a lysine oxidase that produces hydrogen peroxide (21) and exhibits antibacterial activity against other Gram-negative and Gram-positive bacteria (22). The controlled AlpP-mediated autolysis of subpopulations of cells within the centre of biofilm microcolonies is thought to promote the detachment of dense clusters of cells, regulate biofilm spatial architecture, and facilitate dispersal (8). Although the extracellular biofilm matrix is relatively uncharacterized in *P. tunicata*, a recent study identified and characterized a novel protein, SLR4, as an abundant matrix component (23). SLR4 not only forms the protective S-layer around cells in a planktonic state, but also around extracellular components of the biofilm matrix including outer membrane vesicles and filaments (23).

Despite previous knowledge of biofilm development in *P. tunicata*, there exist hundreds of proteins of unknown function encoded within the *P. tunicata* genome (5), many of which may play important biofilm-related functions. Here, to further elucidate the molecular mechanisms and identify key proteins responsible for biofilm development in *P. tunicata*, we have performed a time-course shotgun proteomic analysis of *P. tunicata* cells from a planktonic to early and late biofilm states. Our analysis reveals hundreds of candidate biofilm-associated proteins, including many hypothetical proteins of unknown function. We then investigated the top scoring biofilm-associated protein, EAR30327 (which we designated as “BapP”), and characterized its function using reverse genetic methods. Our work establishes BapP as a novel Ca^2+^-dependent adhesin required for biofilm formation in *P. tunicata*, thus contributing to our understanding of marine biofilm development and diversity of Bap-like biofilm adhesins in bacteria.

## RESULTS AND DISCUSSION

### Proteomic analysis of biofilm development in P. tunicata

To explore the proteomic determinants of *P. tunicata* biofilm development, we performed comparative liquid chromatography-tandem mass spectrometry (LC-MS/MS) shotgun proteomics of *P. tunicata* D2 liquid cultures grown for 8 h (planktonic), 24 h (biofilm), 48 h (biofilm), and 72 h (biofilm). Pellicle biofilms were grown by culturing cells in liquid flasks in non-shaking (static) conditions (see Methods). We also collected protein from the media at 24-72 h time points and performed proteomics on these samples (Fig. 1). Across all samples, we identified a total of 942 *P. tunicata* proteins with a coverage of at least 1 high-confidence peptide assignment (Table S1). A subset of 288 proteins excluding low abundance proteins was used to generate a heatmap of relative abundance across all samples (Fig. 2A). Proteins clustered into four groups based on their relative abundance profiles: cluster 1 (n = 84) proteins were enriched in media samples, cluster 2 (n = 54) proteins were enriched in planktonic samples, and cluster 3 (n = 69) and 4 (n = 81) proteins were biofilm-associated with cluster 4 proteins showing increased abundance at early biofilm stages (24 h) and cluster 3 proteins showing increased abundance at middle to late biofilm stages (i.e., 48 - 72 h). Planktonic, biofilm, and media samples therefore display unique proteomic profiles. To test this further, we performed principal component analysis (PCA) of all samples based on their proteomic profiles (Fig. 2B). As shown by the PCA, the samples separated distinctly according to their category (planktonic, media, biofilm; Fig. 2B). Biofilm samples also showed some additional separation by time point consistent with the heatmap, but not based on the region of biofilm collected (center versus edge) (Fig. 2B).

**Fig. 1:**
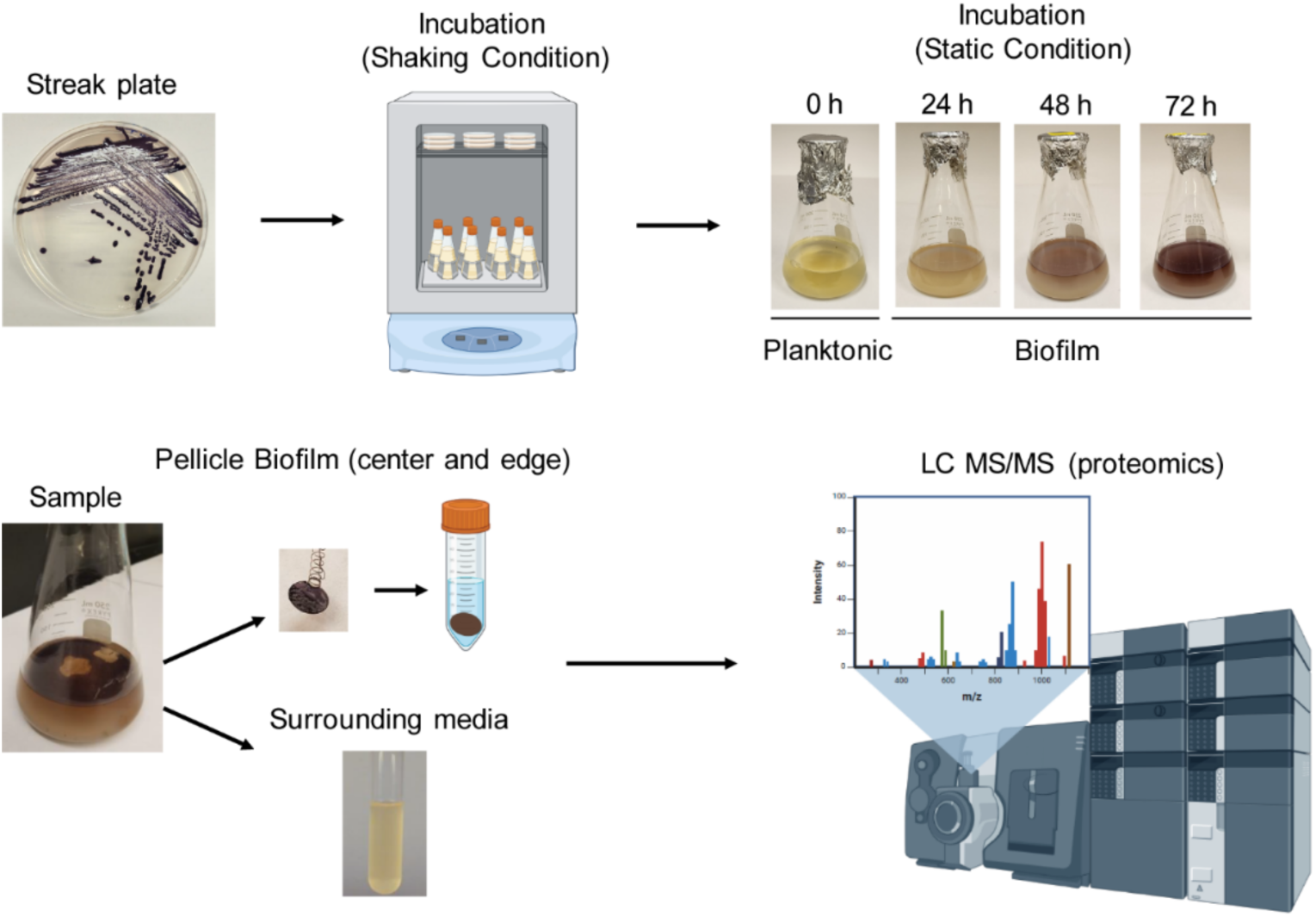
Overview of protocol for sampling of *P. tunicata* pellicle biofilms and planktonic cultures. First, *P. tunicata* was cultured on marine agar streak plates, incubated at 24 °C for 72 h, and used to make liquid cultures in marine broth. In static (non-shaking) conditions, pellicle biofilms began to develop after approximately 24 h and increase in biomass over time at the air-liquid interface. A device was used to extract circular segments from the pellicle biofilm, which was resuspended in Tris-HCl pH 8.3, and used for LC-MS/MS proteomic analysis. For comparison, samples of liquid media below the biofilm surface were also taken for proteomic analysis.

**Fig. 2:**
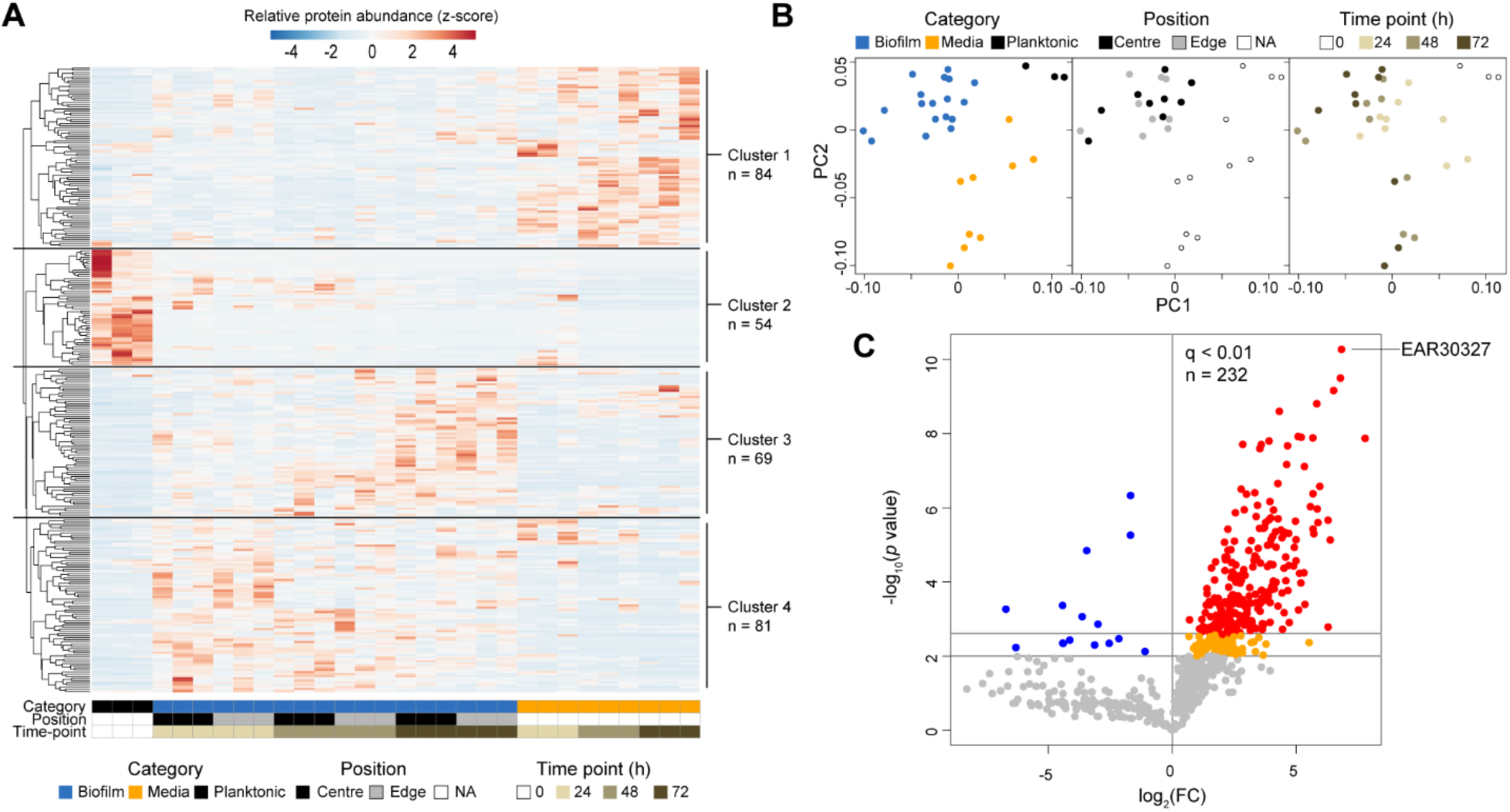
Comparative proteomic analysis of *Pseudoalteromonas tunicata* throughout biofilm development. (**A**) Proteomic abundance heatmap of 288 proteins detected by LC-MS/MS analysis. Per-protein abundances (# spectral hits) were row-normalized to Z-scores depicting relative abundance across samples. Proteins clustered into four groups based on their relative abundance profiles across samples. Cluster 1 proteins are media-enriched, cluster 2 proteins are planktonic-enriched, and cluster 3 and 4 proteins are biofilm-enriched, with cluster 3 associated with mid-to-late stage biofilms and cluster 3 associated with early stage biofilms. (**B**) Principal component analysis (PCA) plot of samples based on their proteomic profiles. Similar to hierarchical clustering in (a), the PCA plot reveals the grouping of samples based on sample type. (**C**) Volcano plot depicting differentially abundant proteins in biofilm samples (N = 18) versus planktonic (N = 3) samples. Proteins with significantly increased abundance in biofilms are located in the top right quadrant, with the top-scoring proteins (q < 0.01) shown in red. Remaining biofilm-associated proteins with weaker significance (*p* < 0.01) are colored orange. Proteins with significantly increased abundance in planktonic conditions are located in the top left quadrant (colored blue). Non-significant proteins are shown in gray.

While the heatmap provides a visual overview of protein abundance profiles, differential abundance analysis was required to detect statistically significant differences. We therefore compared normalized protein abundance across biofilm and non-biofilm conditions to detect putative biofilm-associated proteins (Fig. 2C). Out of the 942 detected proteins, a total of 232 were identified as biofilm-associated based on significant normalized abundance increases (log_2_FC > 0.5 and *q* < 0.01, two-tailed *t*-test, BH adjustment of *p* values) in biofilm versus planktonic samples (Fig. 2C, Table S2). Only 15 proteins were detected with increased abundance in planktonic conditions (Fig. 2C, Table 1), which may be due to lower overall proteomic coverage and fewer biological replicates (n = 3) of these samples.

**Table 1.**
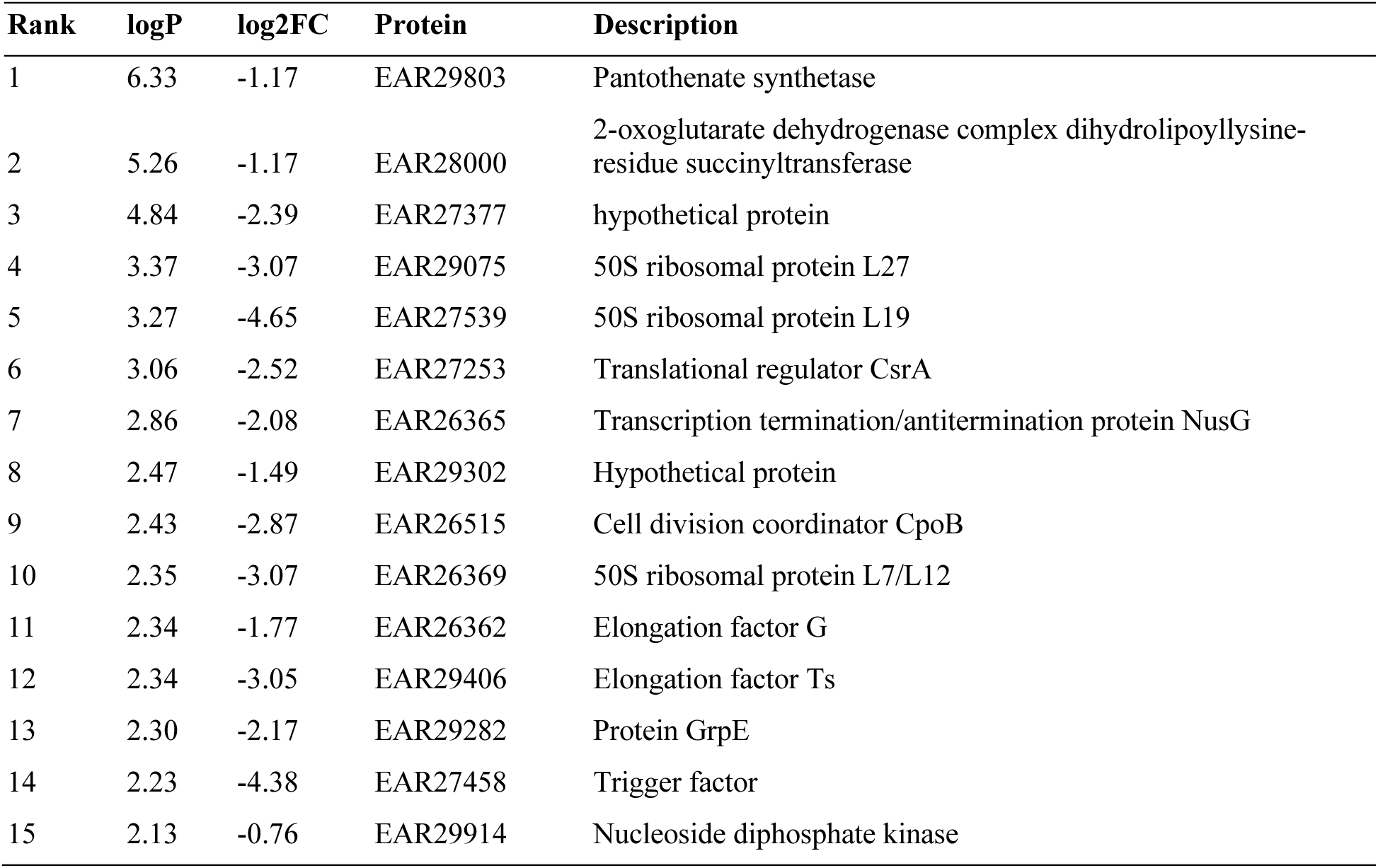
Proteins with increased abundance in planktonic conditions. Descriptions are based on annotations of numerous identical proteins collected from NCBI’s “Identical Protein Groups” resource.

| Rank | logP | log2FC | Protein | Description |
| --- | --- | --- | --- | --- |
| 1 | 6.33 | -1.17 | EAR29803 | Pantothenate synthetase |
| 2 | 5.26 | -1.17 | EAR28000 | 2-oxoglutarate dehydrogenase complex dihydrolipoyllysine-residue succinyltransferase |
| 3 | 4.84 | -2.39 | EAR27377 | hypothetical protein |
| 4 | 3.37 | -3.07 | EAR29075 | 50S ribosomal protein L27 |
| 5 | 3.27 | -4.65 | EAR27539 | 50S ribosomal protein L19 |
| 6 | 3.06 | -2.52 | EAR27253 | Translational regulator CsrA |
| 7 | 2.86 | -2.08 | EAR26365 | Transcription termination/antitermination protein NusG |
| 8 | 2.47 | -1.49 | EAR29302 | Hypothetical protein |
| 9 | 2.43 | -2.87 | EAR26515 | Cell division coordinator CpoB |
| 10 | 2.35 | -3.07 | EAR26369 | 50S ribosomal protein L7/L12 |
| 11 | 2.34 | -1.77 | EAR26362 | Elongation factor G |
| 12 | 2.34 | -3.05 | EAR29406 | Elongation factor Ts |
| 13 | 2.30 | -2.17 | EAR29282 | Protein GrpE |
| 14 | 2.23 | -4.38 | EAR27458 | Trigger factor |
| 15 | 2.13 | -0.76 | EAR29914 | Nucleoside diphosphate kinase |

Planktonic-associated proteins included proteins involved in transcription (e.g., NusG), translation (ribosomal proteins, EF-Tu, CsrA), and cell division (e.g., CpoB). These proteins reflect core physiological processes of intracellular proteins and indicate differences in the metabolic state of cells. CsrA for example is a major regulator of biofilm formation by controlling the expression of biofilm genes and c-di-GMP metabolism (24). LC-MS/MS abundance profiles of example planktonic-associated proteins (EF-Tu and Ribosomal protein L3) are shown in Fig. 3. We also detected twenty proteins with significant abundance increases in media samples (Table S3). Media-enriched proteins may also reflect the activities of planktonic cells or non-adherent cells and, potentially, dispersed cell populations. Several proteins were detected at high levels in 24 h media samples as well as in planktonic cells (e.g., GroEL chaperone, Fig. 3).

**Fig. 3.**
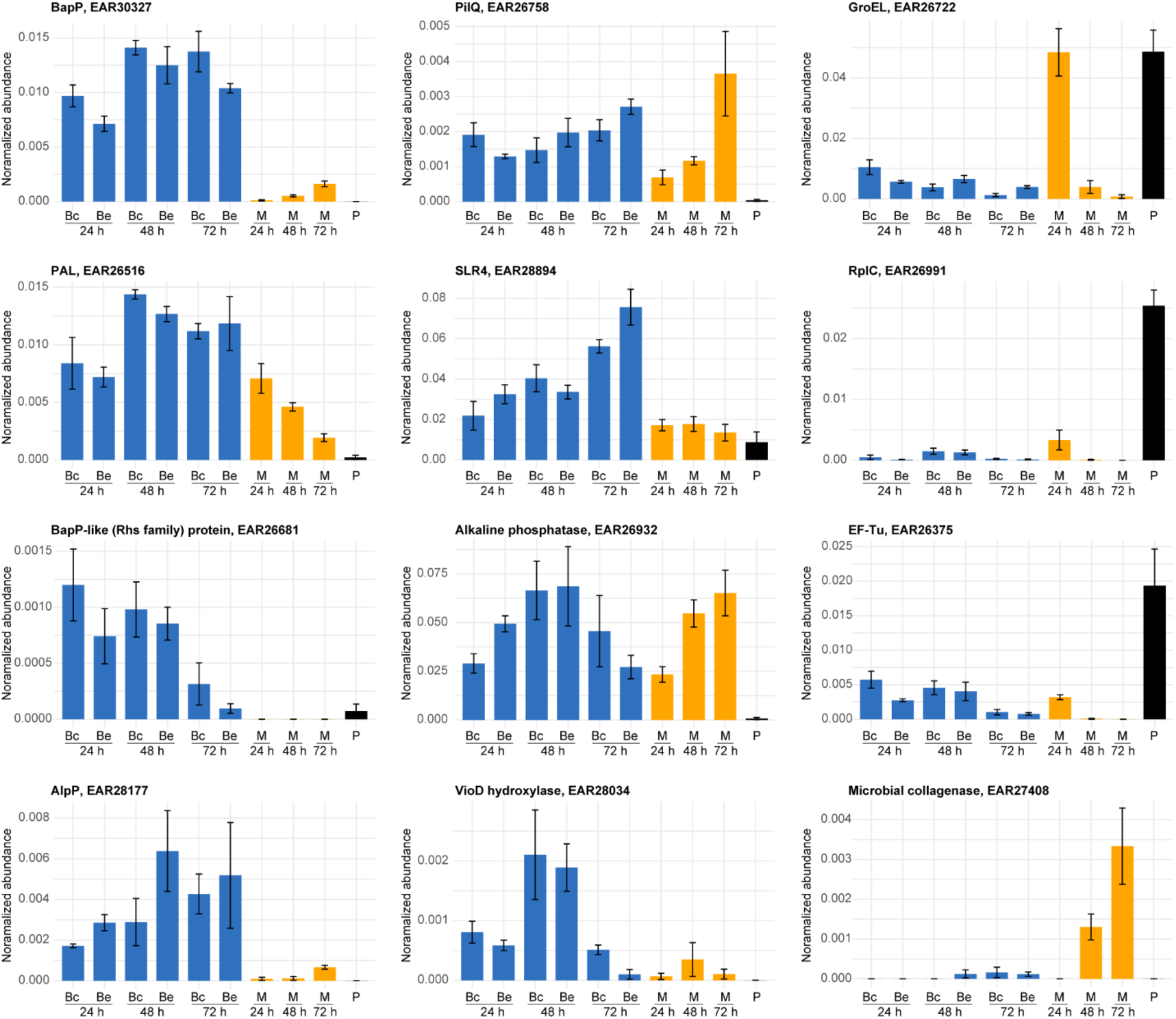
Relative abundance of selected proteins across planktonic, biofilm, and media samples based on LC-MS/MS proteomic analysis. Protein abundance was calculated as a percentage of total peptide spectral matches to the *P. tunicata* proteome. Bc – Biofilm (centre region); Be – Biofilm (edge region); M – media; P – planktonic. Bars represent the mean ± standard error (SE).

### Analysis of biofilm-associated proteins

We first explored the top identified biofilm-associated proteins based on *p* value (top 15 listed in Table 2), which include known biofilm-related proteins as well as proteins with yet-to-be identified roles. The top-ranked protein overall was NCBI/GenBank accession # EAR30327 (Fig. 2C, Table 2), a hypothetical protein that we targeted for subsequent experimental characterization. EAR30327 was detected at high relative abundance in all biofilm time points and reduced abundance in the media and planktonic stages (Fig. 3). The second ranked biofilm-associated protein was PAL (peptidoglycan-associated lipoprotein), a protein known for its role in stabilizing the outer membrane by interacting with the Tol-Pal system (25). The PAL protein showed a similar abundance profile to EAR30327, but in media samples was found at higher abundance at earlier (24 h) time points (Fig. 3). Although the Tol-Pal system has been shown to be up-regulated during biofilm development in other organisms (25, 26), the considerable abundance of PAL in *P. tunicata* biofilms suggests an important, yet-to-be identified role. One possibility is a role related to the formation or structure of outer membrane vesicles which are abundant in the biofilm matrix (23). The type IV pilus secretin PilQ (ranked #5) was also significantly increased in biofilm samples and media samples (Fig. 3). Type IV pili contribute to surface binding and sensing during biofilm formation and may also be involved in uptake of extracellular compounds and DNA (27). The chaperone DnaJ (ranked #7) was increased in biofilm development, consistent with a previous study demonstrating that *E. coli ΔdnaJ* mutants are deficient in biofilm formation (28). Ranked #8 was a 2’ 3’-cyclic nucleotide 2’-phosphodiesterase/3’-nucleotidase, and its increased abundance in biofilm may reflect a putative role of nucleotidases in hydrolyzing extracellular DNA for nutrition (29). Ranked #13 is a predicted peptidyl-prolyl cis trans isomerase (EAR27747). The related enzyme Ppi-B has been shown to contribute to biofilm formation in *Mycobacterium tuberculosis* (30).

**Table 2.** Proteins with increased abundance in biofilm. Descriptions are based on annotations of numerous identical proteins collected from NCBI’s “Identical Protein Groups” resource.

| Rank | logP | log2(FC) | Protein | Description |
| --- | --- | --- | --- | --- |
| 1 | 10.27 | 6.83 | EAR30327 | Ig-like domain-containing protein, "BapP" (this study) |
| 2 | 9.50 | 6.79 | EAR26516 | Peptidoglycan-associated protein |
| 3 | 9.16 | 6.51 | EAR30706 | RimK family alpha-L-glutamate ligase |
| 4 | 8.81 | 5.83 | EAR27711 | S46 family peptidase |
| 5 | 8.60 | 4.32 | EAR26758 | Putative type IV pilus biogenesis protein PilQ (Cytoplasmic ATPase) |
| 6 | 7.90 | 5.20 | EAR27273 | PQQ-binding-like beta-propeller repeat protein |
| 7 | 7.88 | 5.67 | EAR30476 | Chaperone protein DnaJ |
| 8 | 7.87 | 7.78 | EAR30609 | 2' 3'-cyclic nucleotide 2'-phosphodiesterase/3'-nucleotidase bifunctional periplasmic protein |
| 9 | 7.80 | 3.91 | EAR27464 | SurA N-terminal domain-containing protein |
| 10 | 7.67 | 4.65 | EAR28309 | nitroreductase family protein |
| 11 | 7.92 | 5.08 | EAR26410 | FKBP-type peptidyl-prolyl cis-trans isomerase |
| 12 | 7.60 | 3.53 | EAR28471 | Pyrimidine/purine nucleoside phosphorylase |
| 13 | 7.71 | 2.85 | EAR27747 | Uncharacterized protein |
| 14 | 7.11 | 5.33 | EAR26625 | Uncharacterized protein |
| 15 | 6.58 | 5.95 | EAR28769 | Uncharacterized protein |

Out of the top 15, some notable biofilm-associated proteins were two alkaline phosphatases (EAR26932, EAR26933) that were particularly abundant in biofilm samples (Table S1 and S2). Alkaline phosphatases have been shown to be critical for regulating biofilm formation in other organisms (e.g., *P. aeruginosa*) under phosphate depletion stress (31). Ranked #26 was the flagellar hook protein, FlgE, which has been shown to be essential for biofilm formation (32). Other flagellar proteins, including two flagellins (EAR29565, EAR29563), were also enriched in biofilm samples as well as being detected in the surrounding media. Detection of flagellar proteins was expected given the role of motility in initial biofilm formation, intra-biofilm motility, biofilm dispersal, as well as the potential role of flagellins as structural components of the extracellular and biofilm matrix (33).

Among the list of 232 biofilm-associated proteins, we also detected several proteins previously shown to be important for biofilm development in *P. tunicata*. For example, the MSHA pilin protein MshA had increased biofilm abundance, and the violacein pathway proteins (VioA, C, D, and E) were also significantly increased. The tryptophan hydroxylase (VioD) protein showed a unique temporal pattern of expression, because it was at higher abundance at the middle biofilm (48 h) time point (Fig. 3). The production of such pigments in *P. tunicata* is correlated with antifouling activity (34) and has been observed in mature biofilms (7). The autolytic protein, AlpP, was also detected at significantly higher abundance in biofilms, consistent with previous literature (7, 8). AlpP relative abundance increased over time in biofilms, reaching a maximum level in 72 h biofilms (Fig. 3). This finding is consistent with its role in programmed cell lysis to enable cell redistribution and dispersal (8), which likely increases during later stages of biofilm development. The S-layer protein, SLR4, also increased in abundance throughout biofilm development from approximately 2% abundance in 24 h biofilms to 7.5% abundance in 72 h biofilms (Fig. 3), consistent with our previous study which identified SLR4 as a major component of the *P. tunicata* biofilm matrix (23). The shedding of surface-layer proteins such as SLR4 may contribute to biofilm structure and help to form a protective layer around extracellular biofilm matrix components (23, 35). The SLR4 protein was the second most abundant protein detected overall (alkaline phosphatase EAR26932 was #1) and was also significantly enriched in biofilm compared to planktonic samples.

Lastly, among the top biofilm-associated proteins are hypothetical proteins whose relevance to biofilm development is unclear, including EAR27747, EAR26625, and EAR28769, which were among the top 15 biofilm-associated proteins (Table 2), and others beyond this list (e.g., EAR30269). AlphaFold modelling of EAR27747 revealed tandem bacterial immunoglobulin-like (BIg) domains similar to the structure of the invasin and intimin family of adhesins (36). The predicted structure of EAR30269 (AF-A4C3Y7-F1), which was highly abundant in biofilms but also detected in media and planktonic samples, had a beta-helical fold similar to the *S. pneumoniae* surface adhesin, PfbA (37). Thus, in addition to pili, *P. tunicata* appears to produce a variety of surface adhesin proteins involved in biofilm development, most of which are uncharacterized.

### Structural modeling predicts EAR30327 is a Ca2^+^-dependent adhesin

We further investigated the top identified biofilm-associated protein, EAR30327, an uncharacterized protein annotated as “Ig-like domain-containing protein”. BLAST searches of EAR30327 identified homologs in only six additional species (all members of *Pseudoalteromonas*) with >50% coverage and *E-* value < 0.001 (Fig. S1). EAR30327’s closest identified relative was an orthologous protein in *P. ulvae*, an organism also known to colonize the marine host, *Ulva lactuca* (38) (Fig. S1). As all identified EAR30327 homologs were also uncharacterized proteins, EAR30327 is a distinct protein of unknown function that is conserved in multiple species of *Pseudoalteromonas*.

To gain insights into the function of EAR30327, we applied AlphaFold (39) to predict its structure (Fig. 4A). The predicted structure of EAR30327 has an N-terminal domain (residues 21-246) with a 5-bladed β-propeller fold, followed by 12 immunoglobulin-like β-sandwich folds (Fig. 4A). The 12 β-sandwich repeats can be subdivided into three classes of internal repeats (Fig. 4B, 4C). The first six repeats (R1-R6) and a seventh repeat (R7) found closer to the C-terminus, adopt a cadherin-like Greek-key structure that possess matches to the Calx-β PFAM (PF03160) domain. A second class of shorter repeats match the bacterial immunoglobulin (BIg) 9 PFAM family (PF17963) (Fig. 4B, 4C).

**Fig. 4:**
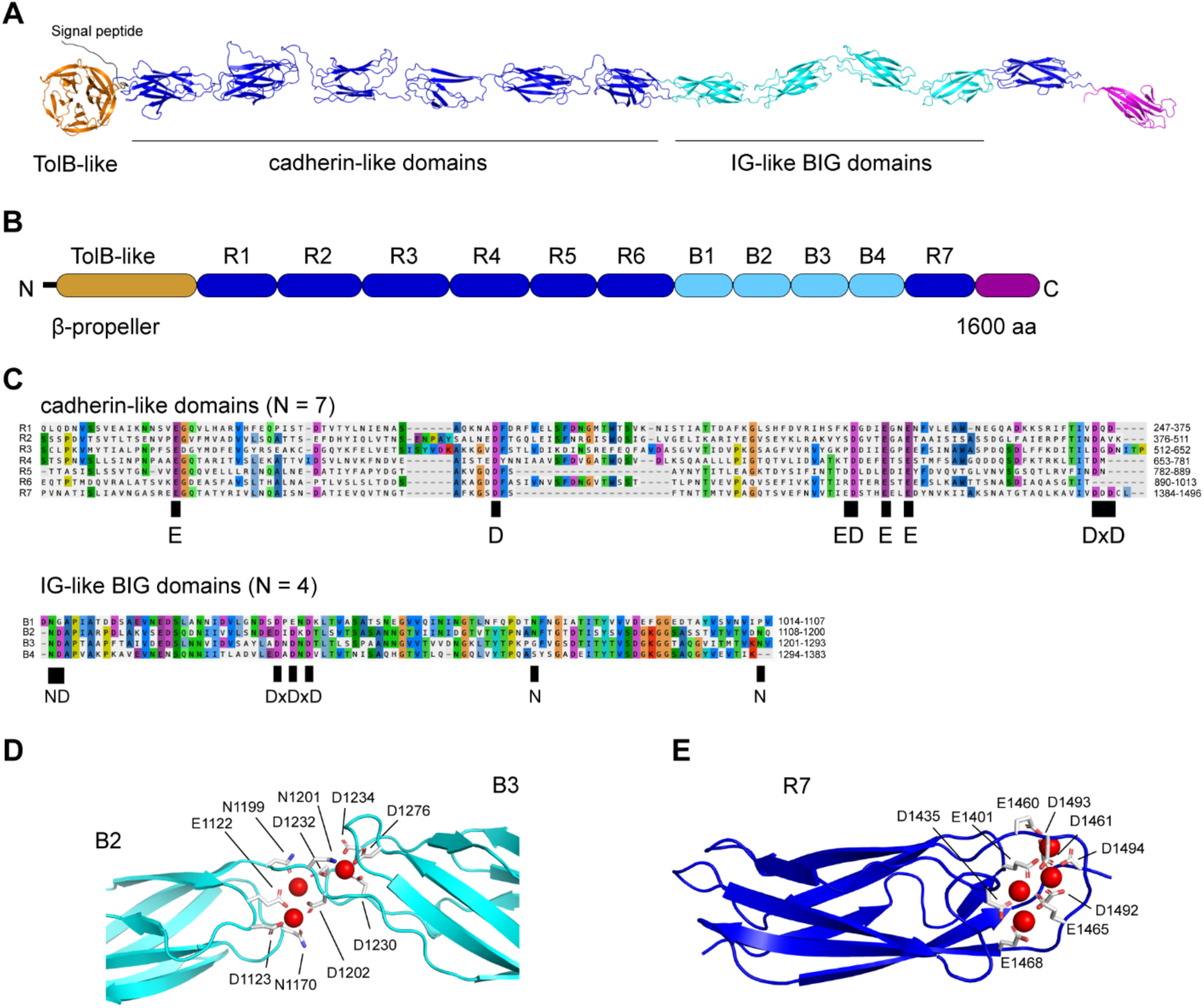
Sequence and structural analysis predicts EAR30327 (“BapP”) as a Ca^2+^-dependent outer membrane adhesin. (**A**) AlphaFold model of BapP, colored by its domain architecture shown in (**B**). The predicted structure of BapP consists of an N-terminal Sec/SPI signal peptide (amino acids 1-20), an N-terminal 5-bladed propeller, and 12 tandem beta-sandwich repeats. The beta-sandwich repeats can be subdivided into three classes based on alignments shown in (**C**). (**D**) and (**E**) show predicted Ca^2+^-binding sites in B2, B3, and R7 domains based on AlphaFold and AlphaFill modeling (see Methods). Key Ca^2+^-binding residues are also depicted in the alignments in (**C**).

The structural architecture of EAR30327 resembles that of other biofilm associated protein (Bap) adhesins (40–42). For instance, tandem cadherin-like domains are present in the giant RTX adhesin, SiiE, from *Salmonella enterica* (43), and also the large, secreted proteins (e.g., CabD and CabC) from the marine bacterium, *Saccharophagus degradans*, which mediates Ca^2+^-dependent homophilic and heterophilic interactions (44) and have carbohydrate-binding activity (45). Calx-β like domains have also been identified in the large RTX adhesin protein, LapA (46). We therefore aligned EAR30327 to these other adhesins to further examine potential similarities. The beta-propeller N-terminal domain of EAR30327 did not align to the N-terminal domains of other known adhesins, and appears to be a unique feature of the BapP. The tandem β-sandwich repeats of EAR30327, however, aligned with similar regions of other adhesins including LapA and CabD, with significant *E*-values and identities exceeding 30% (Fig. S2).

Within the EAR30327 sequence, we identified matches to putative Ca-binding motifs identified previously in bacterial CHDL domains (39), including DxD motifs which were found in both the cadherin-like and BIg repeats (Fig. 4C). Using AlphaFold3 (47) in combination with AlphaFill (48), we modeled EAR30327’s interactions with Ca^2+^ ions (Fig. 4D, E). Numerous Ca^2+^ binding sites were predicted in the linker regions between consecutive β-sandwich domains (Fig. 4D, E), which is a common mechanism also found in Ca^2+^-dependent RTX adhesins (41, 49). The binding of Ca^2+^ ions to similar regions of RTX adhesins is thought to enable rigidification of their extender domains, allowing them to project outward to reach their targets (41, 49).

Thus, together the sequence and structural modeling results strongly suggest a Ca^2+^-binding adhesin function for *P. tunicata* EAR30327. We therefore designated this protein as *Pseudoalteromonas tunicata* biofilm adhesin protein or BapP.

### BapP is required for proper biofilm formation and surface adhesion

To test the predicted function of BapP as a biofilm adhesin, we generated a Δ*bapP* knockout mutant and verified it by PCR (Fig. 5A) and sequencing of amplified PCR product from its genomic DNA (see Methods). A silver stained SDS-PAGE gel of WT supernatants revealed a distinct band that matched BapP by LC-MS/MS analysis, and this band was absent in the Δ*bapP* strain, again confirming successful deletion of the whole *bapP* gene (Fig. 5B). This result also suggests that BapP is at least partially released extracellularly or, alternatively, that it is loosely associated with the cell surface. This is consistent with the biology of non-fimbrial adhesins involved in biofilm formation, which are secreted by the T1SS and loosely attached to the cell surface and often released into the extracellular space (42).

**Fig. 5:**
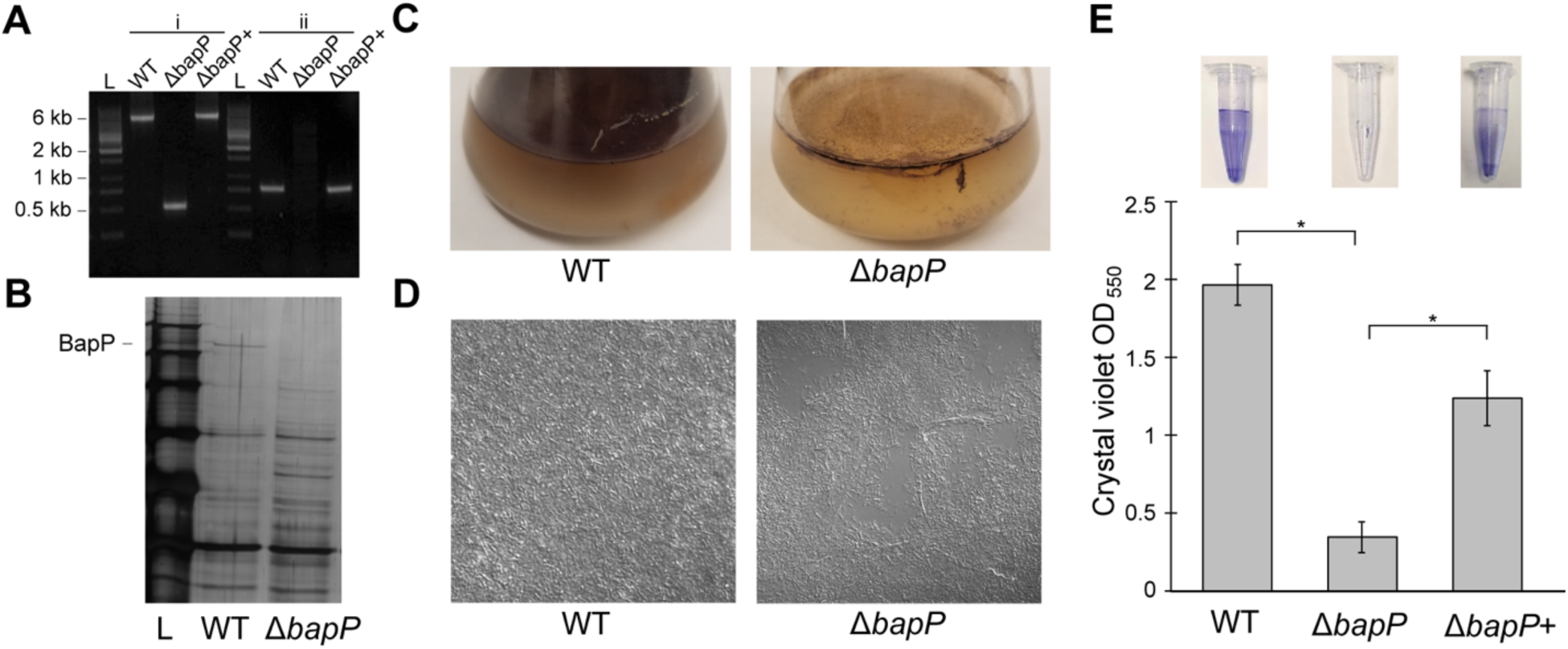
Loss of BapP reduces biofilm formation and surface adhesion. (**A**) Verification of *P. tunicata* strains by PCR amplification. Lanes 2-4 marked “i” are PCR products amplified with primers JC563 and JC564. An amplicon of 510-bp was expected in *ΔbapP* mutant, and a 5301-bp fragment was present in both wild type and *ΔbapP*+ strains. Lanes 6 – 8 marked “ii” are PCR products amplification with primers JC563 and JC565. A 722-bp fragment was present in wild type and *ΔbapP*+ but absent in *ΔbapP* strain due to absence of JC565 binding site. (**B**) Silver stain SDS-PAGE gel of supernatant containing extracellular proteins. A unique band was observed in WT supernatants and missing in Δ*bapP*. The band was excised from the gel and identified as BapP by LC-MS/MS. (**C**) Pellicle biofilms of WT versus Δ*bapP* formed by 72-hour incubation in marine broth under static conditions. (**D**) Confocal microscopy of 72-hrs pellicle biofilms (63 X magnification). (**E**) Crystal violet assays of WT, Δ*bapP*, and rescue (Δ*bapP*+) strains cultured for 24 h in centrifuge tubes containing marine broth. 95% confidence intervals are shown as error bars. The asterisks indicate significant differences in biofilm formation between WT and Δ*bapP* and between Δ*bapP* and Δ*bapP+* (*p*_adj_ < 0.01, two tailed t-test, Bonferroni correction, 15 replicates per condition).

We cultured pellicle biofilms and examined their morphology over a 24-72 h period. As seen after 72 h, the WT formed robust pellicle biofilms with high cell density, whereas the Δ*bapP* mutant formed thin fragile biofilms with a loss of cohesion between cells (Fig. 5C). Regions of the Δ*bapP* biofilm were also visibly detached from the glass substrate. Similarly, confocal microscopy revealed a dense and homogeneous layer of cells in WT biofilms, whereas Δ*bapP* had a fragmented and heterogeneous biofilm with a loss of cell-cell cohesion (Fig. 5D).

To examine the reduced biofilm-forming capability of the Δ*bapP* mutant quantitatively, we performed crystal violet biofilm assays using centrifuge tubes. The WT formed robust biofilms on these tubes within a 24 h period, reaching a maximum OD_550_ of 2.0 units, whereas there was a significant (5.7-fold) reduction in biofilm formed in the Δ*bapP* mutant (*p* = 8.6 × 10^-19^; Fig. 5E). To verify the genotypes, we generated a rescue mutant by replacing the in-frame Δ*bapP* (ATGGGATCCTAA) with the wildtype *bapP* gene in the genome of the Δ*bapP* strain, which was then verified by PCR amplification (Fig. 5A). The Δ*bapP*+ rescue mutant successfully restored biofilm formation in the crystal violet assay (Fig. 5E). The BapP band also reappeared in the supernatant of the Δ*bapP*+ strain confirming successful mutant rescue (Fig. S3). We also confirmed the identity of this band as BapP by mass-spectrometry analysis.

### Ca***^2+^***-dependent BapP-mediated biofilm formation

Given the presence of numerous putative Ca^2+^-binding sites in BapP (Fig. 4) and the established role of Ca^2+^ in RTX biofilm adhesins (41, 49), we investigated the influence of Ca^2+^ on biofilm formation in the WT versus Δ*bapP* strain. To control the CaCl_2_ concentration, we used non-marine complex media without added Calcium (“CM1”) as described in the Methods. Crystal violet assays were then performed with and without the addition of 1.8 g/L CaCl_2_ (16.2 mM) added to the CM1 media, which is equivalent to the Calcium concentration of Difco marine broth 2216. In the WT strain, we observed significant increases in biofilm formation in the presence of added CaCl_2_, with a pronounced effect at later time points (Fig. 6). At the 72-h time point, crystal violet OD_550_ readings increased from a mean of 1.5 units (no added CaCl_2_) to a mean of 4.4 units (1.8 g/L added CaCl_2_) (*p*_adj_ = 1.8×10^-5^) (Fig. 6). By comparison, in the Δ*bapP* strain the addition of Ca^2+^ only increased OD_550_ readings from 0.6 units to 1.7 units (N.S., *p*_adj_ > 0.01) (Fig. 6). The weak residual Ca^2+^-dependent increase in biofilm formation in the Δ*bapP* mutant may be due to the activities of additional Ca^2+^-dependent biofilm adhesins in *P. tunicata* beyond BapP (e.g., EAR26681, which shows partial similarity to the repetitive region of BapP, Fig. S4).

**Fig. 6:**
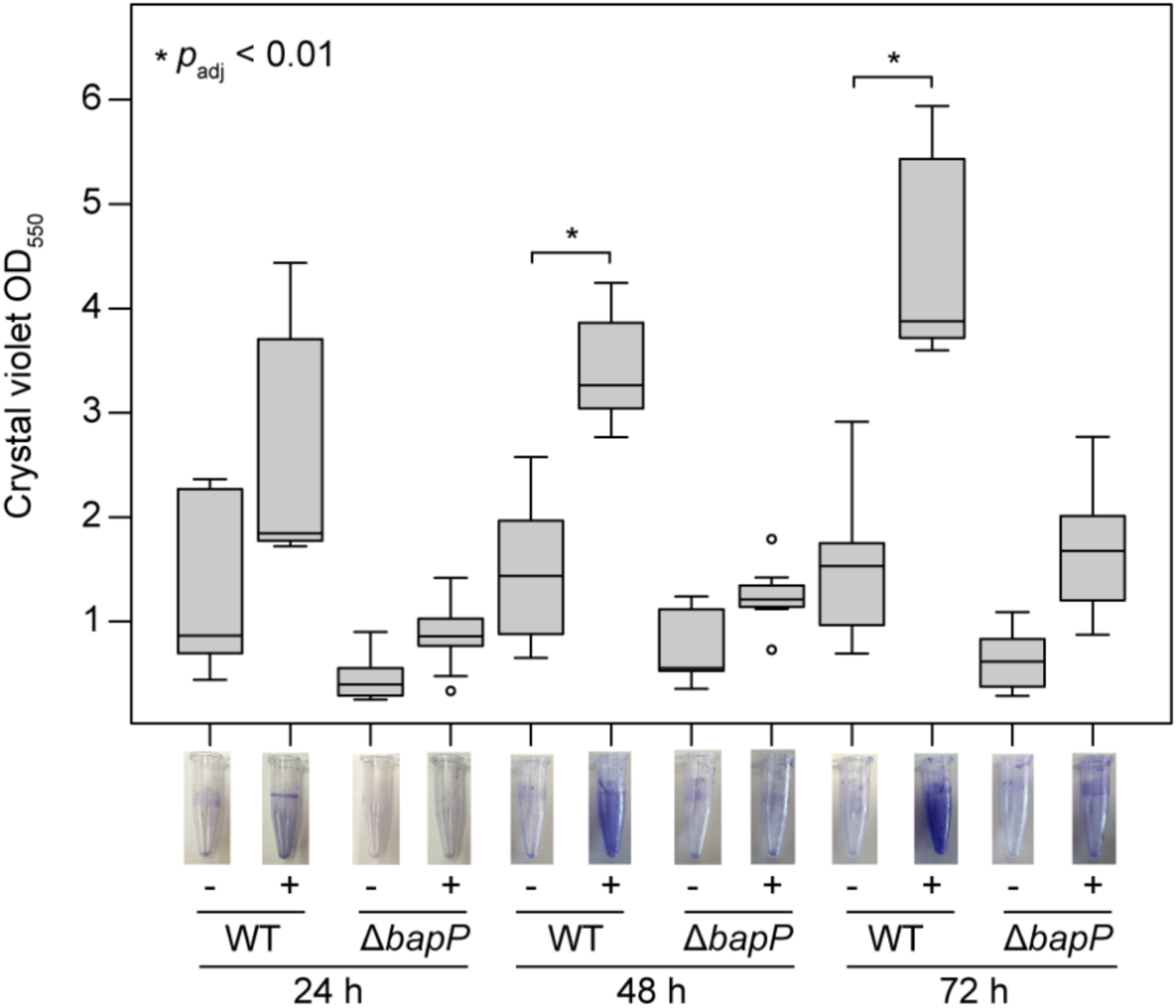
Quantification of the effect of Ca^2+^ addition on biofilm formation. Crystal violet biofilm assays were performed by culturing WT and Δ*bapP* cells in CM1 media over 24, 48, and 72 h periods, with (+) and without (-) the addition of 1.8 g/L (16.2 mM) of CaCl_2_. The boxplots depict the distributions of OD_550_ readings of cells attached to the walls of centrifuge tubes (6 replicates per sample). In the WT strain, when BapP is present, the addition of Ca^2+^ is associated with a significant (*p*_adj_ < 0.01, two-tailed t-test with Bonferroni correction) increase in biofilm levels, but this is not observed in the Δ*bapP*. Boxplots show the lower quartile, median, and upper quartile of the data, with whiskers extending to 1.5 times the interquartile range (IQR) above the third quartile or below the first quartile. Images of representative tubes are shown for each condition.

To account for the possibility that Δ*bapP* had a growth defect, we measured the OD_600_ of all cells in the centrifuge tubes including the media (Fig. S5). Despite the WT showing increased biofilm formation based on crystal violet assays, the total concentration of cells in the tubes including the media was higher for Δ*bapP* than that of the WT. This indicates that the observed effects (Fig. 6) are not due to growth artifacts, and that the majority of Δ*bapP* cells remained in solution, non-attached to the substrate.

### The BapP gene cluster encodes a putative Type 1 secretion system

To gain further insights into the function of BapP, we examined its genomic neighborhood and used AlphaFold3 to structurally annotate adjacent protein-coding genes (Fig. 7). Immediately downstream of *bapP*, we identified four components of a T1SS including a membrane fusion protein (MFS, EAR30323), a LolD-like inner membrane ATPase subunit (EAR30322), and tandem genes encoding ABC transporters (EAR30321 and EAR30320) (Fig. 7A). Using AlphaFold3’s multimer modeling, we predicted the complex formed by these components (Fig. 7B, Fig. S6). Remarkably, the predicted EAR30320-EAR30322 complex resembles the general structure of tripartite efflux systems (AcrAB and MacAB) and the T1SS-associated ABC transporter complex HlyBD, all of which export substrates through the TolC outer membrane pore. Fig. 7B depicts our model of a secretion system formed by the EAR30320-EAR30323 complex interacting with the TolC protein of *P. tunicata* (EAR29116).

**Fig. 7:**
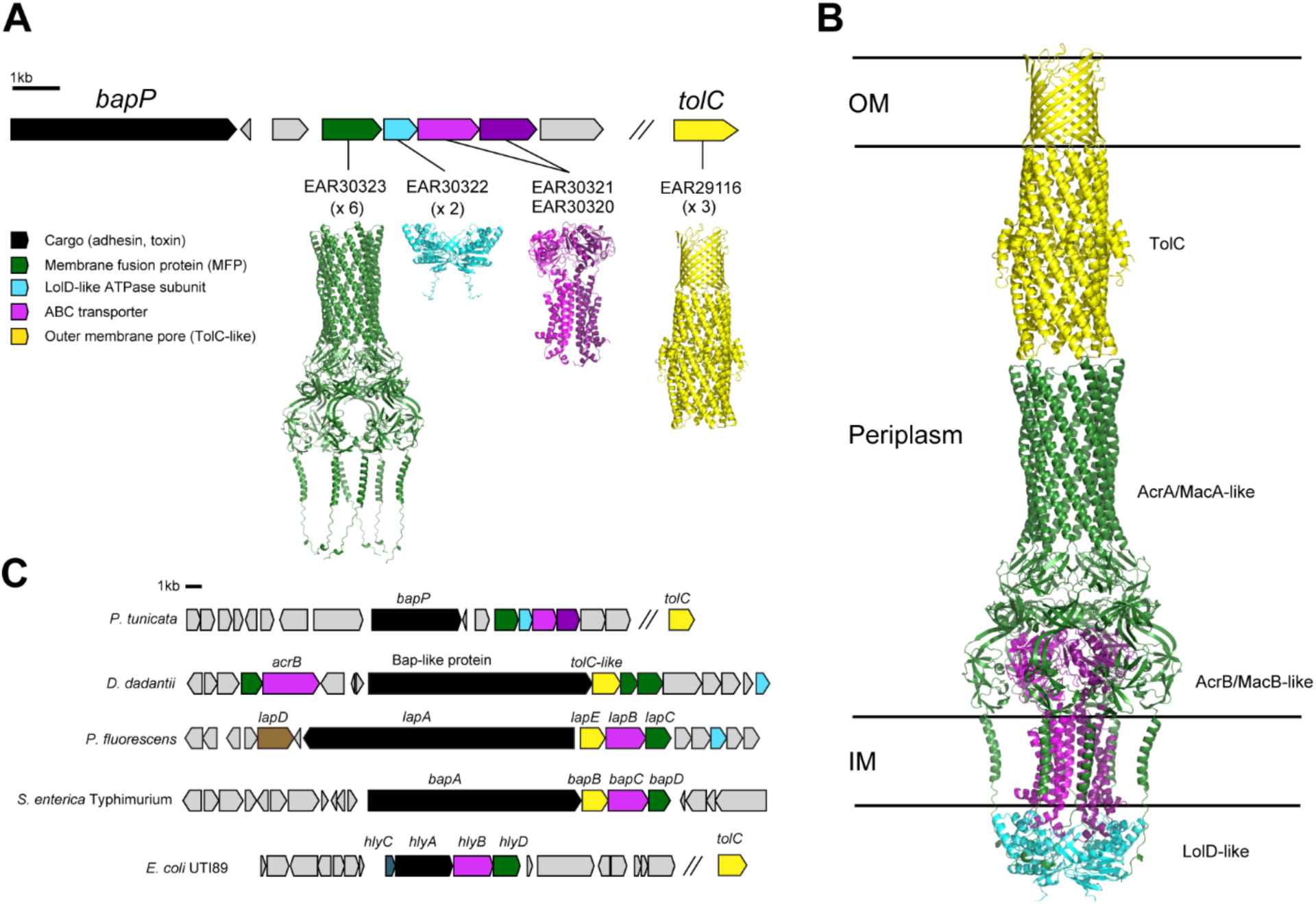
BapP is encoded next to a putative Type 1 secretion system. (**A**) *bapP* gene cluster and AlphaFold3-predicted structures of proteins encoded by the downstream genes, EAR30320-EAR30323. (**B**) AlphaFold3-predicted structure of the protein complex formed by EAR30320-EAR30323 placed alongside the predicted structure of TolC (EAR29116). EAR30320-EAR30323 were predicted as a membrane fusion protein (MFP), ABC transporter, and LolD-like ATPase subunit. As shown in (**C**), genes encoding these components are commonly found in other T1SS-secreted adhesins and toxins, along with TolC in some cases. The genomic context similarities and structure predictions strongly suggest that BapP is encoded next to, and thus likely secreted by, a T1SS formed by these components.

Figure 7C shows the gene neighborhoods surrounding other T1SS-secreted adhesins and toxins, including *lapA* from *P. fluorescens*, *bapA* from *S. enterica*, hemolysin A (*hlyA*) from *E. coli*, and a putative adhesin in *D. dadantii* which had detectable homology to the repetitive region of BapP. Similar to BapP, each of the T1SS-secreted proteins are encoded adjacent to a MFP and ABC transporter which form a T1SS with a TolC-like pore. TolC-like genes are found immediately adjacent to P. fluorescens lapA, S. enterica bapA, and the Bap-like gene in D. dadantii, but they are encoded distally in *E. coli* and in *P. tunicata*.

Together, the sequence and structural modeling of BapP’s surrounding gene neighborhood strongly suggests a T1SS secretion mechanism. This, combined with other evidence presented (sequence, structural, and functional) indicates that BapP and related proteins form a novel, distinct family of T1SS-secreted biofilm adhesins, which are new members of a broad and diverse class of Ca_2+_-dependent Bap-like adhesins in bacteria.

## CONCLUSION

Biofilm-formation by marine bacteria is critical for marine microbial ecology and also has substantial practical, industrial and economic implications (50). In this work, we explored the proteomic determinants of biofilm development in the marine microorganism *P. tunicata*, which serves ecologically important roles in marine biofilms and possesses a mostly uncharacterized proteome containing an abundance of hypothetical proteins. Comparative shotgun proteomics detected hundreds of proteins with increased abundance in *P. tunicata* biofilms, many of which are either uncharacterized or have currently unknown roles in biofilm development. Longitudinal analysis over a 24-72 h period also revealed temporal patterns of protein expression, reflecting changes in biofilm function in different stages of biofilm development. Through in-depth proteomic analysis of biofilms, we were able to overcome the technical challenges of separating proteins from what can be highly complex and challenging samples to work with.

In addition to identifying hundreds of biofilm-associated proteins and providing a unique hypothesis-generating proteomic dataset for future studies, our work led to the discovery of a Ca^2+^-dependent adhesin (BapP) in *P. tunicata* and related species, which we confirmed experimentally as playing a key role in biofilm formation. BapP appears to share structural and functional similarities with other biofilm adhesin proteins (e.g., LapA), but is distantly related in sequence and also possesses entirely unique domains (a predicted N-terminal beta-propeller). The BapP gene cluster also suggests a T1SS mechanism similar to other RTX adhesins, establishing BapP as a new member of the broad class of Bap adhesins. Additional work is required to characterize the regulation and localization of BapP, as well as the relative contribution of BapP along with other putative adhesins produced by *P. tunicata*. Knockout studies of different segments of BapP would also be helpful in determining the function of its N-terminal domain, and the different classes of beta-sandwich repeats in its structure.

## METHODS

### Culturing of *P. tunicata* samples throughout planktonic to biofilm development

A frozen stock of *P. tunicata* strain D2 was streaked on Difco marine agar 2216 and incubated for 48 h at room temperature. An overnight culture was made by inoculating a colony into 4 mL of marine broth, which was incubated overnight at 24 °C with 120 rpm. Eight subcultures were made in Erlenmeyer flasks by transferring 1 mL (OD_600_ = 1.3) of the overnight culture into 100 mL of marine broth. The flasks were incubated for 8 h at 24 °C with gentle shaking (100 rpm). After 8 h incubation, two of these cultures were used as planktonic cultures and considered “T0” samples. The six remaining flasks were used to generate pellicle (air-liquid interface) biofilms, by incubating these flasks statically at room temperature for 3 days. Samples were collected at 24 h, 48 h, and 72 h of incubation. Duplicate samples were collected from each time point, and they were pooled in one tube. Biofilm samples were collected from the center and the edge of the biofilm in addition to media samples. The entire procedure was repeated three times to provide biological replicates. Thus, a total of 30 samples were collected from four different time points (T) as follows: T0 = [(planktonic 8 h shaking) (n = 3)]; T24 = [24 Biofilm sample center (n = 3), Biofilm sample edge (n = 3), media (n = 3)]; T48 = [48 Biofilm sample centre (n = 3), Biofilm sample edge (n = 3), media (n = 3)]; T72 = [72 Biofilm sample center (n = 3), Biofilm sample edge (n = 3), media (n = 3)].

A method used by Park et al. (51) with some modifications was followed for sample processing. To collect planktonic samples, 7.5 mL was aspirated from each of two flasks using a sterile serological pipette, yielding a total volume of 15 mL. The planktonic cultures were then concentrated to a volume of 2.5–3 mL using a Vivaspin 20 centrifugal filtration unit with a 3 kDa cutoff, operated in a swing bucket at 4000 x *g* for 140–160 minutes at 4 °C. The concentrate (2.5–3 mL) was washed using the same Vivaspin 20 unit with 10 mL of cold Tris-HCl buffer (0.1 M, pH 8.3). This washing step was performed with a fixed-angle rotor at 6000 x *g* for 10 minutes at 4 °C. A final volume of 500 µl of concentrate was collected, and the samples were then frozen at −20 °C.

Pellicle biofilm samples were collected using a clean round coverslip (Fisher brand, No. 2 - 0.25 mm thick, size 18 mm) attached to a custom-designed tool composed of a spring and metal rod shown in Fig. S7, which was cleaned with water and alcohol after each use. Biofilm samples (18 mm diameter) were collected from the edge and the biofilm’s center. The coverslips containing biofilm were placed in a clean tube containing 5 mL of cold Tris-HCl buffer (0.1 M, pH 8.3). Biofilm samples were washed off the surface of the coverslip tube by vortexing gently until the biofilm was completely separated from the coverslip and transferred to clean tubes. The biofilm suspension was centrifuged once at 12,000 x *g* for 10 min at 4 °C. The supernatant was discarded. The pellet was collected and resuspended in 500 µl of cold Tris-HCl (pH 8.3) and vortexed until all the pellet was resuspended. The samples were frozen at −20 °C. Media samples were also collected from each biofilm flask after the biofilm samples were taken and prepared using the same protocol as described above for planktonic samples.

### Shotgun proteomics

#### Sample processing for LC-MS/MS

Prior to LS-MS/MS analysis, samples were processed for protein extraction and quantification. Three rounds of freeze/thaw cycles were performed using 1 L of liquid nitrogen for 30 seconds followed by transfer to a room temperature water bath. Samples (990 µl) were then treated with 10 µl of protease inhibitor cocktail FOCUS ProteaseArrest (G-Biosciences) and kept on ice. A 3 mm sonicator tip (Qsonica Sonicator) was used to sonicate the samples, which were placed in 2 mL round-bottom Eppendorf tubes. Sonication was performed in four cycles on ice, with each cycle consisting of 30 seconds at 30% amplitude, followed by 60 seconds of cooling time between sessions. The sonicated samples were centrifuged (6000 x *g*, 10 min, 4 °C) to remove cellular debris, and the supernatants were collected. The samples were stored at −20 °C for further processing. To perform LC-MS/MS, at least 50 µg of protein sample was collected for each sample. The protein concentration in the lysates was measured using the Bradford assay (52).

#### Proteomics

Ten µg samples were reduced with 10mM dithiothreitol (Sigma-Aldrich, Missouri, USA) alkylated with 20mM iodoacetamide (Sigma-Aldrich, Missouri, USA), acetone precipitated and digested overnight with MS grade trypsin (Promega, Wisconsin, USA). Digested samples were lyophilized then resuspended in 0.2% Trifluoroacetic acid (TFA) and desalted using a homemade C18 zip tip (resin: Empore, 2215-C18(Octadecyl)). The C18 desalted samples were resuspended in 18 µl buffer A (0.1% FA buffer pH 2.7). Six microliters of each sample were injected into the *tims*TOF Pro (Bruker Daltronics, Bremen, Germany) using nanoflow liquid chromatography using a Bruker NanoElute chromatography system (Bruker Daltronics, Bremen, Germany). Liquid chromatography was performed at a constant flow of 400 µl/min and a 15cm reversed-phased column with a 75 µm inner diameter, packed with Reprosil C18 (PepSEP, Bruker, Germany). Mobile phase A was 0.1% Formic Acid, and Mobile phase B was 99.9% Acetonitrile, 0.1% Formic Acid. The *tims*TOF Pro was outfitted with a CaptiveSpray source (Bruker Daltronics, Bremen, Germany), operated in PASEF mode. MS and MS/MS scans were limited to 100 m/z to 1700 m/z, and a polygon filter was applied to the m/z and ion mobility dimensions to select for multiple charged ions most likely to be peptide precursors. Collision energy was applied as a function of ion mobility, using a linear regression with the following parameter settings: 0.85 V·s/cm² at 27 eV and 1.30 V·s/cm² at 45 eV. TIMS voltage was calibrated using ions from the Agilent Tune Mix (m/z 622, 922, 1222). Active exclusion of MS/MS scans was enabled with a setting of 0.40 min. Quadrupole isolation was set to 2 m/z for ions with m/z less than 700, and 3.0 m/z for ions with m/z greater than 800. All MS experiments were completed at the Bioinformatics Solutions Inc. MS lab (Waterloo, Ontario, Canada).

MS Raw Files were processed using PEAKS XPro (v10.6, Bioinformatics Solutions Inc., Ontario, Canada). The data were searched against a custom database containing the *P. tunicata* proteome. Precursor ion mass error tolerance was set to 20 ppm and fragment ion mass error tolerance was set to 0.05 Da. Semi-specific cleavage with trypsin was selected with a maximum of 2 missed cleavages. A fixed modification of carbamidomethylation (+57.02 Da) on cysteine residues was specified. Variable modifications of deamidation (+0.98 Da) on asparagine and glutamine, as well as oxidation (15.99 Da) on methionine, were specified. The false discovery rate threshold was set to 1% for the database search.

Normalized protein abundance calculated by Peaks was further converted to proportions of overall counts using R v4.3.3. Heatmaps of protein abundance were generated using pheatmap (https://github.com/raivokolde/pheatmap) with row-normalization applied using the scale=“row” option. PCA was performed using the “prcomp” function in R. Differential analysis of protein abundance was performed using two-tailed t-tests with *p* values adjusted for multiple-hypotheses using p.adjust with the Benjamini-Hochberg method (53). Differentially abundant proteins were identified as those with log_2_FC values > 0.5 and *q* values < 0.01.

### Bioinformatic analysis of EAR30327

#### BLAST and phylogenetic analysis

A comprehensive set of EAR30327 (BapP) homologs were identified by BLAST analysis of 80,789 proteomes from the GTDB R214 database using BapP as a query and an *E*-value threshold of 0.001. The taxonomic distribution of genomes possessing one or more BapP homologs was summarized and visualized using AnnoTree (54). A smaller set of more closely related homologs was identified by a BLAST search of the NCBI nr database on July 1, 2024 using an *E*-value threshold of 0.001 and a query coverage threshold of 50%. A phylogenetic tree of BapP and close homologs from other *Pseudoalteromonas* species was produced using PhyML v3.1 (55) using 1018 sites and the LG model with 4 rate classes.

#### Sequence and structural modeling of EAR30327 and the surrounding gene cluster

Structural modeling of EAR30327 was initially performed using ColabFold’s implementation of AlphaFold2 (39, 56) and later refined using AlphaFold3 (47). Structural models were then edited to obtain a linear domain representation for improved visualization in PyMOL v3.0 (https://pymol.org/), and connective loops closed within MODELLER (57). Internal sequence repeats based on the structural model were further aligned using MUSCLE (58) and further edited and visualized using AliView (59). To allocate metal-binding sites, AlphaFill (48) was used to identify potential Ca^2+^ binding sites based on homology to other Ca^2+^-binding template structures, and identified Ca-binding domains were then submitted individually or as pairs to AlphaFold3 for placing high-confidence calcium ions (pLDDT > 75) as in context of the full-length protein sequence only low-confidence sites could be predicted (pLDDT < 25).

Gene neighborhoods surrounding BapP and other selected proteins (T1SS-secreted adhesins and toxins) were visualized and explored using AnnoView (60). Structural and functional annotation was performed using a combination of AlphaFold3 (47) predictions and existing sequence-based annotations (CDD, Interpro) from the NCBI and Uniprot databases. Based on the stoichiometry of components in AcrAB-TolC and MacAB-TolC systems, an AlphaFold3 model was generated for a protein complex composed of six monomers of the putative membrane fusion protein EAR30323, two monomers of the LolD-like subunit EAR30322, and a heterodimer of the ABC transporter proteins EAR30320 and EAR30321. The resulting complex was then manually placed alongside the predicted trimer structure of *P. tunicata* TolC (EAR29116).

### Construction of Δ*bapP* and rescue (Δ*bapP*+) plasmids

#### Bacterial strains, plasmids, and growth media

Bacterial strains and plasmids used in this study are derived from earlier work (4, 61, 62) and are listed in Table S4. *E*. *coli* strains were grown at 37 °C in LB medium (1.0% tryptone, 0.5% yeast extract and 0.5% NaCl). *P. tunicata* was grown in Difco marine broth 2216. Enzymes were obtained from New England Biolabs (NEB). Antibiotics were used at the following final concentrations: kanamycin, 50 μg/ml for *E. coli* or 100 μg/ml for *P. tunicata*; chloramphenicol, 20 μg/ml.

#### Plasmid construction

Q5 high-fidelity DNA polymerase (NEB) was used for plasmid construction and confirming constructs. The upstream region (1,469 bp) of *bapP* gene and 23-bp downstream of the gene were PCR amplified using oligos JC559 and JC560 (Table S5, Fig. S8) and its downstream region (1,690 bp), 20-bp upstream of the ORF and start code (ATG) were obtained using primers JC561 and JC562. Both DNA fragments were gel purified and pooled in equal amount as PCR template using JC559 and JC562. The PCR product was digested with EcoRI-XbaI and then inserted into the same sites in pK19mobsacB, yielding plasmid pJC296.

A DNA fragment (6818 bp) containing *bapP* ORF, upstream and downstream regions (989 bp and 1,014 bp respectively) of the gene were PCR amplified using oligos JC679 and JC680 (Table S5, Fig. S8). After gel purification, the amplicon was restricted with EcoRI-XabI and then cloned into the same sites in pK19mobsacB, to obtain plasmid pJC302. The plasmid constructions were verified by restriction enzyme mapping.

#### Construction of P. tunicata strains

Plasmid pJC296 was conjugated into *P. tunicata* D2 with the helper plasmid pRK600 (63). Single cross-over recombination of the pJC296 *into P. tunicata* genome was selected on marine agar with kanamycin. A single colony was streak purified on a fresh plate. A colony was grown in Difco marine broth 2216, diluted serially, and plated on marine agar plates containing 5% sucrose. Resulting clones were tested for kanamycin sensitivity (double cross-over and loss of plasmid backbone). Genomic DNA was isolated from the Kan^S^ clones. DNA fragments were PCR amplified using primer pairs of JC563-JC565 and JC563-JC564. The products were resolved with 1.5% TAE agarose gel. The amplicon generated with oligos JC563 and 564 was also Sanger sequenced at The Centre for Applied Genomics (Toronto, Ontario).

In order to rescue *bapP* ORF in the in-frame-deletion Δ*bapP* mutant, plasmid pJC302 was transferred into the 11*bapP* mutant strain with the plasmid pRK600. Following selection of single cross-over recombination on marine agar with kanamycin, one transconjugant clone was streak purified on a new selection plate. A Kan^R^ colony was then incubated overnight in marine medium, diluted serially, and plated on marine agar with 5% sucrose. Resulting clones were screened for kanamycin sensitivity (*bapP* ORF inserted and loss of plasmid backbone). In order to detect the replacement of in-frame 11*bapP* ORF in mutant, Genomic DNA was extracted, and PCR amplification was performed using primer pairs of JC563-JC565 and JC563-JC564.

### Crystal Violet Biofilm Assays

A modified protocol based on a previous study (64) was used to perform crystal violet assays. WT, Δ*bapP*, and rescue (Δ*bapP*+) strains were grown overnight in Difco marine broth 2216 at 24 °C with shaking at 120 rpm. Overnight cultures were diluted to an OD600 of 0.01 with Difco marine broth 2216, and 900 µL of each subculture was dispensed into regular retention 1.5 mL microcentrifuge tubes (GeneBio). The centrifuge tubes were incubated statically at room temperature for 24 h. Five replicates were made for each sample, and the entire experiment was replicated three times to ensure reproducibility. After the appropriate incubation time, the culture was discarded. The biofilm adhered to the walls of the microcentrifuge tubes was stained by adding 950 µl of 0.1% crystal violet to each microcentrifuge tube and left to stain for 15 min. Then the stain was disposed, and the excess stain was removed by shaking out the tube, and slowly washing with water three times. The trapped water was shaken out, the biofilm was left to dry overnight. To quantify the biofilm, the concentration of crystal violet was measured. 1.1 mL of 30% acetic acid was added to each microcentrifuge tube to solubilize the crystal violet for 15 min. 1.1 mL of 30% acetic acid was added to three microcentrifuge tube to serve as a blank. The absorbance was quantified using a spectrophotometer (BioSpectrometer, Eppendorf) at 550 nm.

The same procedure described above was then used to investigate the effects of CaCl₂ on BapP biofilm formation with one modification: the overnight cultures of WT and Δ*bapP* strains were grown in Complex Media “CM1” broth (tryptone 10 g, yeast extract 5 g, NaCl 10 g, MgSO_4_ 0.150 mg, deionized water 1 L) with and without 1.8 g/L (16.2 mM) CaCl_2_. Each culture was diluted to an OD_600_ of 0.01 with a volume of the corresponding CM1 broth. Then each subculture was dispensed into microcentrifuge tubes. Three replicates were made from each subculture, and the entire experiment was replicated three times to ensure reproducibility.

### Confocal Laser Scanning Microscopy

Pellicle biofilm samples were collected using a clean square coverslip (Fisher brand, No. 1.5 - 0.17 mm thick, Size 18 x 18 mm) and the same custom-designed tool that was described earlier to collect the biofilm samples for proteomic analysis. The coverslip was immediately inverted on a clean microscopic slide and the edges of the coverslip were sealed with a transparent nail polish to avoid biofilm dryness. Slides were examined within one hour of preparation using a Zeiss LSM 700 confocal microscope with the differential interference contrast (DIC) II condenser setting at magnifications of 40X and 63X.

### SDS-PAGE

WT, Δ*bapP*, and Δ*bapP^+^* strains were grown overnight in 4 mL Difco marine broth 2216 at 24 °C with shaking at 120 rpm. Overnight cultures were subcultured, by first standardizing the overnight to an OD_600_ of 1.0 with Difco marine broth 2216. The diluted overnight cultures were then subcultured to 1:99 dilution into fresh marine broth. A total volume of 4 mL was prepared and incubated for 5 hours at 24 °C with shaking at 120 rpm. Each culture was centrifuged at 5000 x *g* and 4 °C for 15 min. The supernatant was transferred to a new container and centrifuged again under the same conditions. The resulting supernatant was treated with two different protease inhibitors to prevent protein degradation: 10 µl of FOCUS ProteaseArrest added to 990 µl of the sample, and 10 µL/mL of 0.5 M EDTA. The samples were stored at −20 °C. On the following day, samples were subjected to an SDS-PAGE (10% SDS) for protein separation based on molecular weight. A 25 µl of samples were loaded into the gel. In addition, 5 µl of BLUelf prestained ladder with MW 5-245 kDa (FroggaBio) was loaded. The gel was run at 90 kV for 20 min, and 140 kV for 60 min. The SDS-PAGE gel was stained using silver staining (Pierce silver stain for Mass Spectrometry, Thermo Scientific) following the kit protocol.

## Author Contributions

S.A., A.S., B.J., S.G., G.C., and J.C., performed experiments. S.A., A.S., G.C., H.T., X.W. U.E., and A.C.D. performed data analysis. T.C.C. and S.E. provided bacterial strains and resources. S.A., A.S., U.E., J.D.N., T.C.C., and A.C.D. interpreted data and designed experiments. All authors contributed to writing and approved the final manuscript.

## Acknowledgements

We thank Dr. Barbara Moffatt for use of her laboratory for culturing of *P. tunicata* pellicle biofilms. This work was supported by an Ontario Early Researcher Award and an NSERC Discovery Grant (RGPIN2019-04266) awarded to A.C.D.

## Competing Interests

All authors declare no financial or non-financial competing interests.

## Data Availability

Data is provided within the manuscript or supplementary information files. Additional raw datasets used and/or analyzed during the current study are available from the corresponding author on reasonable request.

## Notes

### Competing Interest Statement

The authors have declared no competing interest.

