## Supplementary Figures for "Comparative proteomics of biofilm development in *Pseudoalteromonas tunicata* discovers a distinct family of Ca^2+^-dependent adhesins"

**SUPPLEMENTARY MATERIAL**

A

| Description | Total Score | Query Cover | E value | Per. ident | Acc. Len | Accession |
|---|---|---|---|---|---|---|
| Ig-like domain-containing protein [Pseudoalteromonas tunicata] | 3231 | 100% | 0 | 100 | 1600 | WP_009836625.1 |
| Ig-like domain-containing protein [Pseudoalteromonas tunicata] | 3226 | 100% | 0 | 99.81 | 1600 | WP_305975243.1 |
| Ig-like domain-containing protein [Pseudoalteromonas tunicata] | 3221 | 100% | 0 | 99.69 | 1600 | WP_306107941.1 |
| Ig-like domain-containing protein [Pseudoalteromonas ulvae] | 2479 | 100% | 0 | 75.62 | 1600 | WP_086743690.1 |
| Ig-like domain-containing protein [Pseudoalteromonas ulvae] | 2477 | 100% | 0 | 75.62 | 1600 | WP_193331482.1 |
| Ig-like domain-containing protein [Pseudoalteromonas spongiae] | 1640 | 86% | 3e-100 | 29.46 | 2341 | WP_336436765.1 |
| Ig-like domain-containing protein [Pseudoalteromonas spongiae] | 966 | 86% | 4e-96 | 28.51 | 2341 | WP_100914984.1 |
| Ig-like domain-containing protein [Pseudoalteromonas sp. T1lg24] | 791 | 70% | 2e-92 | 28.85 | 1660 | WP_105171922.1 |
| Ig-like domain-containing protein [Pseudoalteromonas piratica] | 952 | 87% | 1e-91 | 29.17 | 2341 | WP_040136050.1 |
| Ig-like domain-containing protein [Pseudoalteromonas sp. P1-9] | 955 | 86% | 1e-89 | 28.35 | 2341 | WP_054980746.1 |
| Ig-like domain-containing protein [Pseudoalteromonas spongiae] | 953 | 86% | 1e-89 | 28.58 | 2341 | WP_010559390.1 |
| Ig-like domain-containing protein [Pseudoalteromonas sp. MMG024] | 952 | 86% | 4e-89 | 28.71 | 2341 | WP_237129147.1 |
| Ig-like domain-containing protein [Pseudomonadota bacterium] | 938 | 86% | 9e-89 | 28.78 | 2341 | MEC8327087.1 |
| Ig-like domain-containing protein [Pseudoalteromonas sp.] | 704 | 72% | 9e-78 | 27.88 | 1538 | WP_372769392.1 |

B

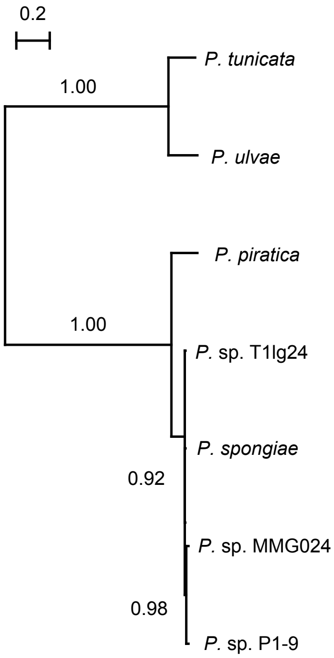


**Figure S1. Identification and phylogenetic analysis of EAR30327 homologs in six additional species of *Pseudoalteromonas*.** A) BLAST result using EAR30327 as a query against the full NCBI nr database. All homologs with E-values < 0.001 and query coverage > 50% are shown. B) Maximum-likelihood phylogenetic tree of EAR30327 from *P. tunicata* along with non-redundant homologs in other *Pseudoalteromonas* species. Bootstrap values are shown above the nodes.

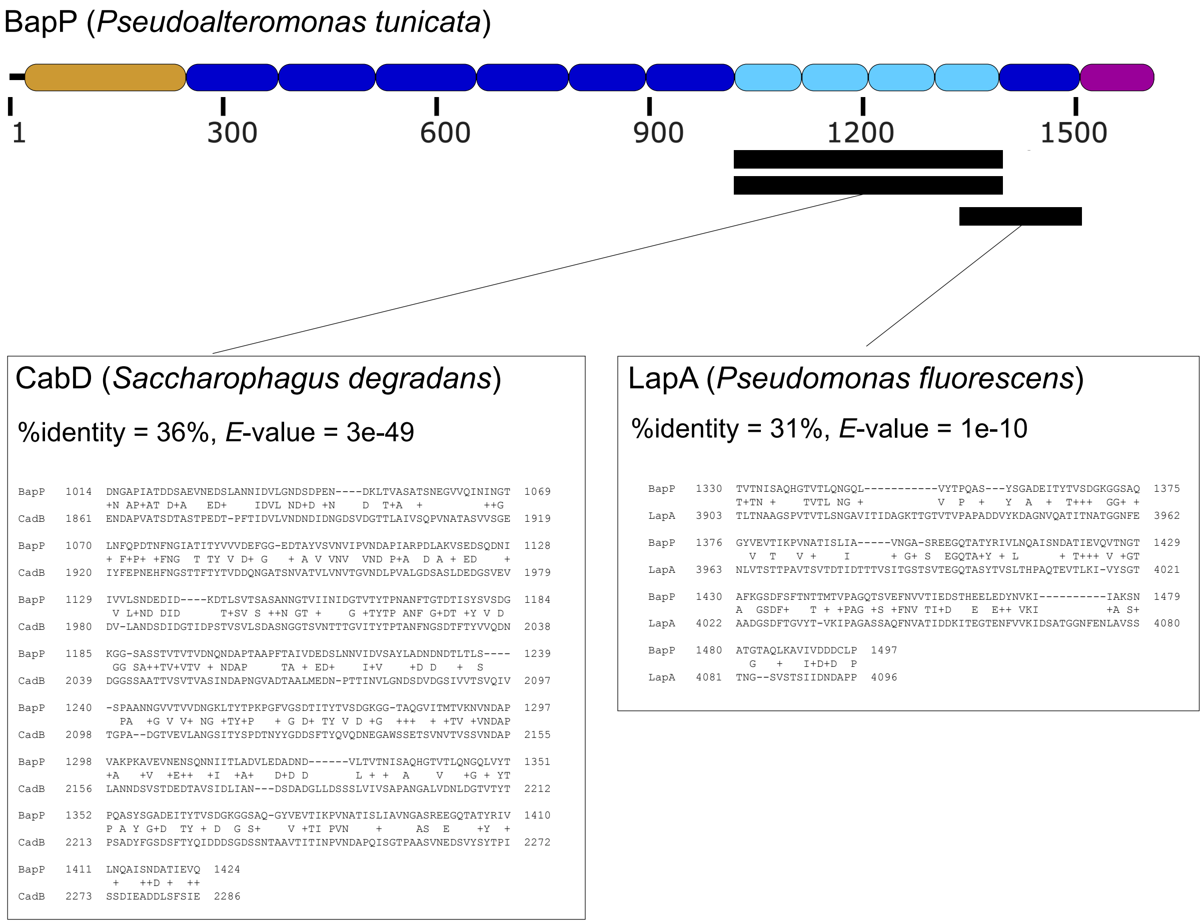


**Figure S2. Partial sequence homology between a putative binding region in BapP and binding domains from other biofilm adhesins.** A domain architectural model of BapP is depicted above, and two regions corresponding to significant BLAST alignments are shown below, the first of which is similar to a region from the CadB adhesin from *S. degradans*, and the second of which is similar to a region from *P. fluorescens* LapA.

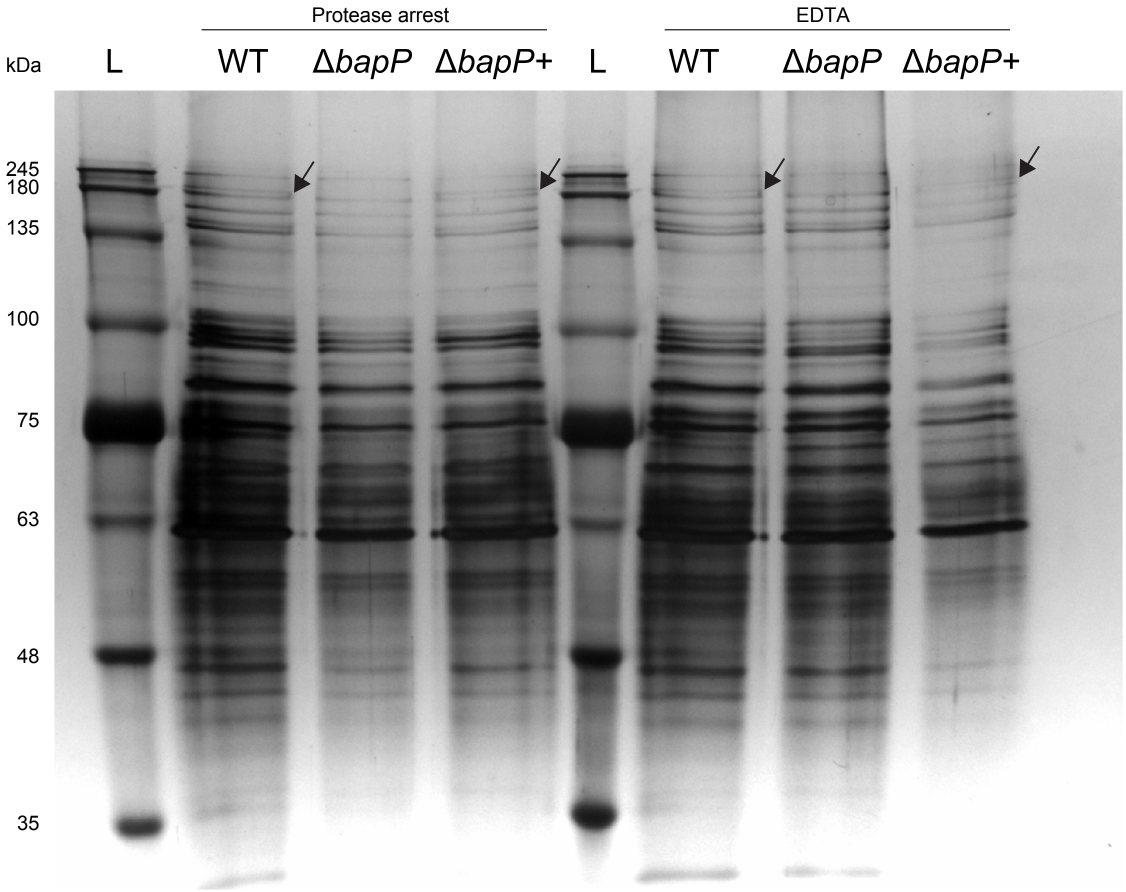


**Figure S3. SDS-PAGE gel of supernatant protein collected from the WT, Δ*bapP*, and Δ*bapP*+ strains.** Protease arrest and EDTA were added to minimize protein degradation. A unique band (arrow) appeared at ~180 kDa in the WT and Δ*bapP*+ strains, and not in the Δ*bapP* deletion strain, and was identified as BapP by LC-MS/MS proteomics.

A

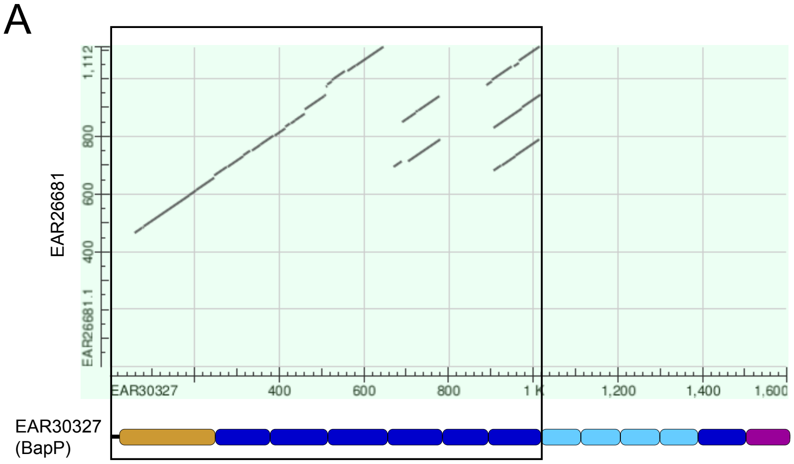


B

```
Score           Expect  Identities     Positives      Gaps
199 bits(506)   1e-55   190/664(29%)   307/664(46%)   95/664(14%)

Query  59     SDALAYQESTDRLYYVTKPVNG--KPRLVYVDMATEQHVDVAATTGTYRLAFSPDGQTLW  116
              SDA+AY +  DRLYY+TK VNG  KP++ YV+M T + V +A   G YRL FSPD   L+
Sbjct  466    SDAVAYDKINDRLYYITK-VNGALKPKVAYVNMQTGEDVKLAEIDGGYRLVFSPDSSQLF  524

Query  117    GSSEDTVFNINTNDGTVSNKVTLTGFATDADKLWGDIVFINDTLHIVTNKKLFAVDLAAG  176
               SS   +  IN   G V +K++L           GD+VFI D L+IV+   L  VDL
Sbjct  525    ASSGRDIVEINPTTGAVISKISLKSNDESITFGMGDLVFIQDELYIVSQLHLIKVDLVNK  584

Query  177    TVREVGMHNLS-VTGSTIDSLGQLLVSS-NAGNNKTDLYTLDPATAKPSLLSSINYRIND  234
                  +G H ++  TG+ +DS G LL+S  N    +T +Y+++    K + +++++Y IND
Sbjct  585    QATVLGKHGVNGATGAEVDSEGNLLISKINNATAQTTIYSINVTELKATAVATVDYAIND  644

Query  235    LATRTYVQPACT--------QLQDNVSSVEAIKNNSVEGQVLHARVHFEQPISTDTVTYL  286
              LA R +    C          ++ ++ ++E I +  VEG  L A+V F      +    +
Sbjct  645    LALRHFDGKTCNTDPVDPVEPIKSSLVAIELISDRQVEGDDLVAKVMFSG--GEEAQLNI  702

Query  287    NIENASAQKNADFDRFVELSFDNGMTWTS---VKNISTIATTDAFKGLSHFDVRIHSFKD  343
              N+ + +A  +ADF   V +SFD+G TW +    KN S     +A   L    ++IH+  D
Sbjct  703    NLSSNTADIDADFTSKVSVSFDDGQTWLTDIEAKNGSVTVPENAKSAL----IKIHTLTD  758

Query  344    GDIEGNENFVLE-AWNEGQADKKSRIFTIVDQDSSSPDVTSVTLTSENVPEGVFMVADVV  402
               ++E +E+  LE ++    +  KS + TIVD+ +              +P+  F+ A  +
Sbjct  759    TEVENDESLRLEVSFAHDPSVVKSELLTIVDKPTGGDTGGGDAC---EMPKVSFITALSI  815

Query  403    LSQATTSEFDHY------IQLVTNSENPAYSALNEDFTGQL-EISFNRGISWQSIGLVGE  455
               +   T +   +      +Q     + PA    N +F  +L +I   + + + ++  V
Sbjct  816    FNGEYTQDSKAFTYEGGEMQFEVGFDGPA--KCNGNFQFKLNDIETTQRLDYSALVSVSS  873

Query  456    LIKAR---------------IYEGVSEYKLRAKVYSDGVTEGAETAAISISASSDGLFAI  500
              L                   + EG   + +R K  +D   E  E   +S+   SD
Sbjct  874    LTNENWLNNVNVALGSATYAVQEGDEGFIVRVKTLADTTKEKNEVFTLSVWNKSDQSDVK  933

Query  501    ERPFTI-NDAV---------------------------KSC-----LPKVMYTIALP---  524
               +  +I N+A                             SC     +P + +  AL
Sbjct  934    YKDHSIENNAATDVDDGTPGTGGGTPGTGTPGNGNTDPDSCSSEDEIPAMKFITALSVDE  993

Query  525    -NPFS-EDGYMDFEVGYRSEAKCDGQYKFELVETSISYVDKAKKGVDFSTLVDIKDINSR  582
                 F+ E G M++  G+  EA C G + F   +      D   KGVD+ST VDI+  + +
Sbjct  994    GRSFTKEGGKMEYYAGFAKEASCSGTFYFSFAD------DHTTKGVDYSTNVDIQTWD-K  1046

Query  583    EFEQFAVDASGVVTIDVPKGSAGFVVRVYGKPDDIIEGPEEFSINAWASPDQSDLFFKDI  642
              +  Q+ VDA+    ++V KG+AGF + +    DD  E  EE+ ++ W   D+SDL  KD
Sbjct  1047   QPAQYNVDAANGAAVNVVKGTAGFTITLTTLADDEAEEREEYFLHTWRKADKSDLKIKDH  1106

Query  643    TILD  646
              TI+D
Sbjct  1107   TIVD  1110
```

**Figure S4. BLAST alignment dot plot depicting regions of similarity between BapP (x-axis) and a related *P. tunicata* protein (EAR26681).** The region of similarity (boxed) covers the N-terminal domain and first six beta-sandwich repeats.

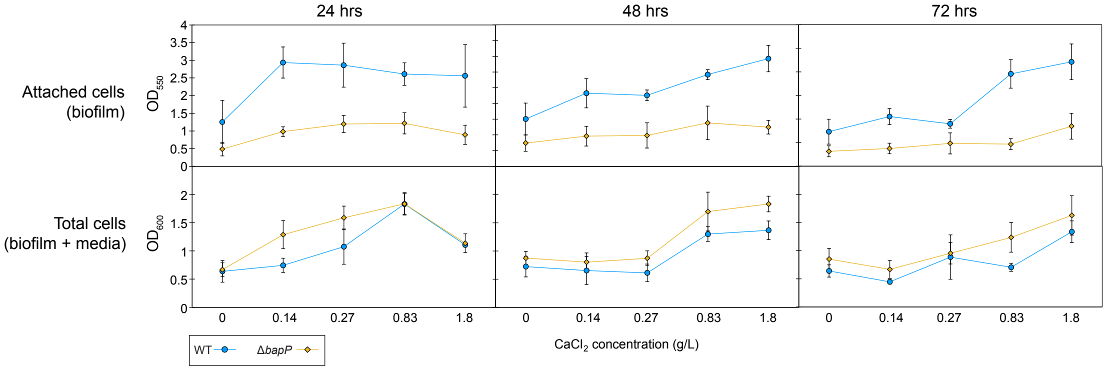


**Figure S5. Line graphs depicting biofilm growth, quantified via 550 nm absorbance, and total cell growth, quantified via 600 nm absorbance, against added $CaCl_2$.** Across all timepoints and $CaCl_2$ levels, the density of WT biofilms exceeds that of Δ*bapP* ($p < 0.05$, two-tailed t-test). However, the total cell growth (attached + non-attached cells) of Δ*bapP* matched or exceeded that of WT cultures. Error bars represent 95% confidence intervals.

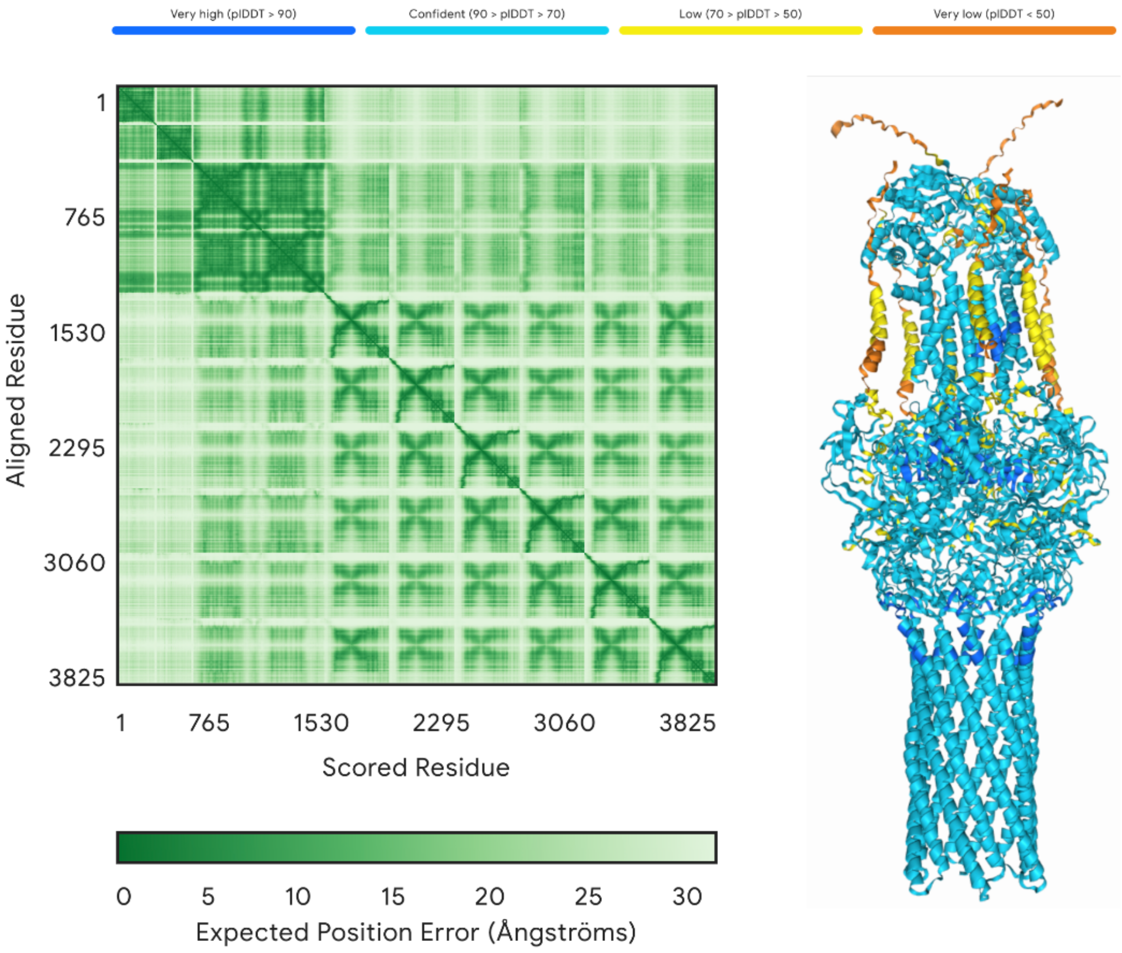


**Figure S6**. **Structural prediction of the EAR30320-EAR30323 complex produced by Alphafold3.** Input sequences included six copies of EAR30323, two copies of EAR30322, and one copy each of EAR30321 and EAR30320. The Alphafold3 heatmap depicting the pairwise interactions between all residues is shown on the left, with the structural model on the right colored by plDDT scores.

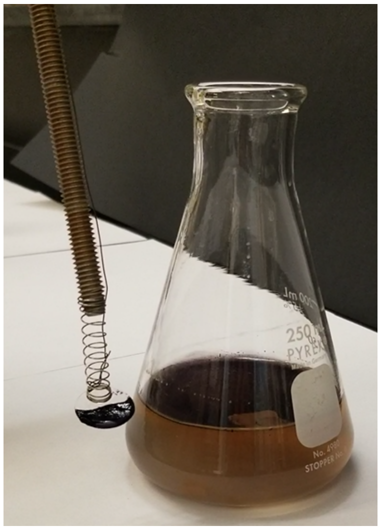

**Figure S7. Biofilm sample collection tool.** A metal dowel was equipped with a tightly coiled spring that holds a glass coverslip. The spring was connected to a piece of fishing line, allowing the coverslip to be carefully inserted beneath the film and then raised to collect a portion of the biofilm. Biofilm samples were collected from the center and edge.

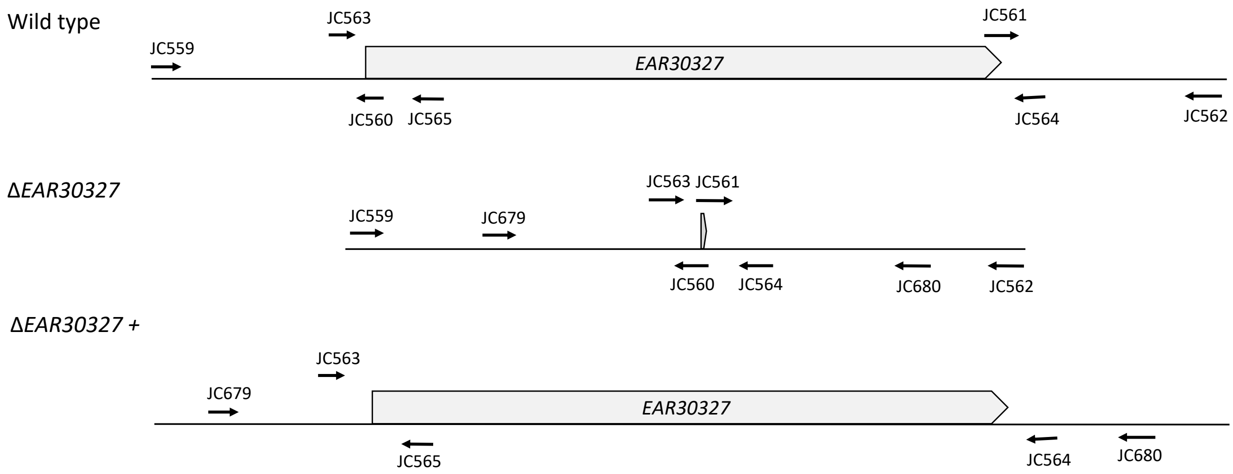


**Figure S8. PCR primer binding regions in *P. tunicata* genomes.**

**Supplementary Table Legends**

Table S1. Proteomics LC-MS/MS protein abundance across all samples. All detected *P. tunicata* proteins are included.

Table S2. Top detected biofilm-associated proteins.

Table S3. Top detected media-enriched proteins.

Table S4. Bacterial strains and plasmids used in this work.

Table S5. Sequences of primers used in this work.
